# Decoupling transcription factor expression and activity enables dimmer-switch gene regulation

**DOI:** 10.1101/2020.04.30.071282

**Authors:** C. Ricci-Tam, J. Wang, I. Ben-Zion, J. Palme, A. Li, Y. Savir, M. Springer

## Abstract

Our mechanistic understanding of how gene regulatory networks can achieve complex biological responses is limited. We find that the galactose-responsive pathway in *S. cerevisiae* has a complex behavior - the decision to induce is controlled independently from the induction level, resembling a mechanical dimmer-switch. Surprisingly, this behavior is not achieved directly through chromatin regulation or combinatorial control at galactose-responsive promoters. Instead, this behavior is achieved by hierarchical regulation of the expression and catalytic activity of a single transcription factor. This genetic motif allows evolution to independently act on both properties, providing a means to tune both resource allocation strategies in dynamic multi-input environments on both physiological and evolutionary time-scales. Hierarchical regulation is ubiquitous and thus dimmer-switch regulation is likely a key feature of many biological systems.

## Main Text

To choose appropriate responses to many different circumstances, cells utilize complex transcriptional programs that integrate multiple inputs from the environment into a complex output. The yeast GAL pathway is a model system for multi-input response (*1, 2*). Here we find that it also exhibits a complex output with two independently controlled output features; the fraction of cells that decide to express the pathway and their expression level. Dissecting this behavior and the underlying molecular mechanism offers a unique opportunity to study complex regulation.

We determined that the complex behavior of the GAL response can be understood in terms of two decoupled responses, a switch-like decision to induce and a rheostat-like (i.e. graded or dimmable) expression level control (Fig. 1, Fig. S1). We measured the steady-state GAL response in 77 different combinations of glucose and galactose by monitoring the output of a *GAL1*pr-YFP, a reporter for expression of the first metabolic enzyme in the GAL pathway (*3*) (Fig. 1A, Fig. S1, Methods). Previously we showed (*1*) that the fraction of cells that decide to induce the pathway (ON fraction) is a one-dimensional function of the galactose:glucose ratio (Fig. 1B,J, Fig. S1C,D). Here we find that the induction level of those ON cells (ON level), in contrast, is well fit by a one-dimensional Michaelian function that depends solely on glucose (Figure 1E,K, Fig. S1E,F, Methods). This design is analogous to a mechanical dimmer-switch controller often used for lights. The galactose:glucose ratio, like the light switch, determines whether cells would turn ON the pathway, i.e. if there is any output. Independently, the glucose concentration, like the light dimmer or rheostat, determines the expression level, i.e. the amount of output.

**Fig. 1.**
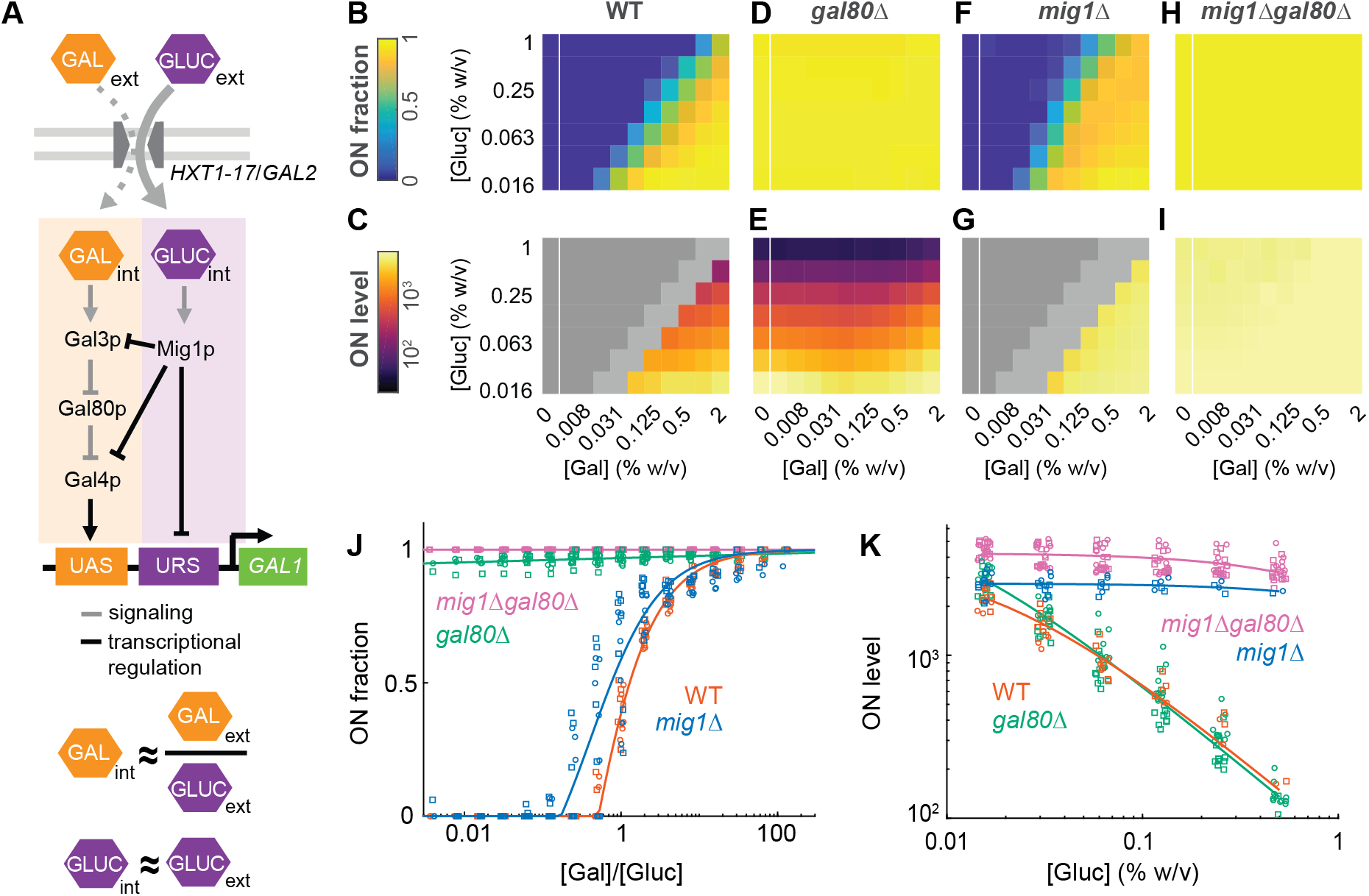
Switch-like decision to induce and rheostat-like control of induction level are genetically decoupled in the yeast GAL pathway. (A) Schematic of the yeast GAL pathway, including hypothesized competitive-transport mechanism (*1*) by which the internal galactose level is a function of the ratio of external sugars, leading the GAL branch of the pathway (orange) to be responsive to the galactose:glucose ratio. (B)-(I), ON fraction and ON level heatmaps of wild type, *mig1*Δ, *gal80*Δ, and *mig1*Δ*gal80*Δ strains in a glucose-galactose double gradient (two replicates for each strain and glucose-galactose combination). (J), A plot of the ON fraction versus galactose:glucose ratio for wildtype, *mig1*Δ, *gal80*Δ, and *mig1*Δ*gal80*Δ strains for experiments in (B)-(I). X-values randomly jittered for visualization purposes. Deletion of *GAL80* eliminates the switch. (K), A plot of the ON level in the ON regime versus the glucose concentration for wildtype, *mig1*Δ, *gal80*Δ, and *mig1*Δ*gal80*Δ strains for experiments in (B)-(I). X-values randomly jittered for visualization purposes. Deletion of *MIG1* eliminates the rheostat. See fit parameters in Table S4.

We find that the switch-and-rheostat response has a genetic basis, where the Gal4p transcriptional activator (*4*) controls the switch and the Mig1p transcriptional repressor (*5, 6*) controls the rheostat (Fig. 1, Fig. S2). We measured the state-state GAL response of a *gal80*Δ, a *mig1*Δ, and *mig1*Δ*gal*80Δ in the same 77 combination of glucose and galactose. The *gal80*Δ ON fraction is always 100%, independently of glucose and galactose, while its ON level responds to glucose concentration similarly to wild type (Fig. 1C,E,K). In contrast, the *mig1Δ* ON fraction depends on the galactose:glucose ratio similarly to wild type, while its ON level is independent of glucose and always at its maximal level (Fig. 1G,K). Thus Gal80p is necessary for the switch and Mig1p is necessary for the rheostat. Consistent with Mig1p and Gal80p being necessary for the rheostat and switch respectively, a *mig1*Δ*gal80*Δ strain is constitutively ON at its maximal level (Fig. 1H-K), i.e. neither ON fraction nor ON level depend on glucose or galactose.

We next sought to determine the molecular mechanism of the switch-and-rheostat response. One possibility is a “chromatin-decoupled regulation” model, inspired by pioneer factors (*7*) and observations in the yeast phosphate (PHO) pathway (*8*), where the switch is controlled by Gal4p-mediated chromatin remodeling and the rheostat by Mig1p-mediated transcriptional regulation (*9*) in fully accessible *GAL1* promoters (Fig. S3A). To test this, we measured chromatin occupancy and gene expression in many glucose and galactose combinations using a modified version of ATAC-seq that incorporates crosslinking and flow-cytometric sorting (*10, 11*) (Fig. S3, Methods). Consistent with our model, the chromatin accessibilities of sorted ON and OFF subpopulations were remarkably different (Fig. 2A, Fig. S3D-F). As expected, the accessibility of the OFF subpopulation appears almost identical to that of maximally repressed wildtype cells. However, contrary to the “chromatin-decoupled regulation” model, which posits that all ON cells would have fully accessible chromatin, we observed a monotonic correlation between ON level and chromatin accessibility at the *GAL1* promoter (Fig. 2A, Fig.S3H-K).

**Fig. 2.**
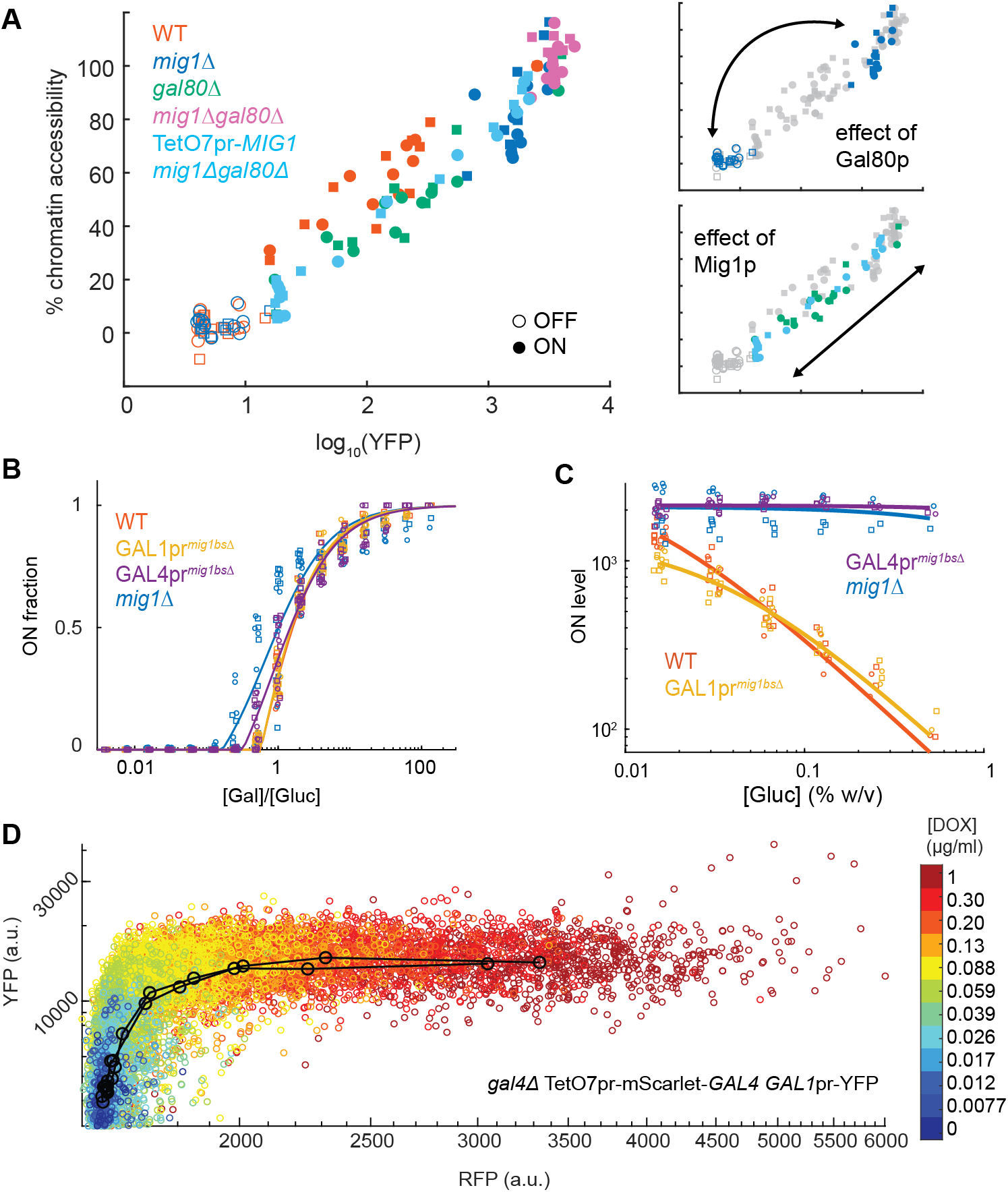
Decoupling occurs through regulation of the Gal4p transcription factor rather than directly at the *GAL1* promoter through chromatin. (A) Scatterplot of YFP expression level (x-axis) versus percent chromatin accessibility (y-axis) at the GAL1 promoter across all samples from wildtype, *mig1*Δ, *gal80*Δ, *mig1*Δ*gal80*Δ, and TetO7pr-*MIG1 mig1*Δ*gal80*Δ strain backgrounds. Data plotted is of two biological replicates (circles indicate replicate 1, squares indicate replicate 2) for each strain and glucose-galactose/doxycycline condition. (B)-(C), ON fraction as a function of the galactose:glucose ratio (B) and ON level as a function of glucose concentration (C) in a glucose-galactose double gradient assay (two replicates) across wildtype, *mig1*Δ, *GAL1*pr^*mig1bsΔ*^, and *GAL4*pr^*mig1bsΔ*^. X-values randomly jittered for visualization purposes. (D) mScarlet versus YFP fluorescence in a *gal4*Δ TetO7pr-mScarlet-*GAL4 GAL1*pr-YFP strain from microscopy of a doxycycline (DOX) titration series (see Fig. S4I for a YFP versus DOX plot). Scatter represents single-cell measurements, with colors corresponding to discrete DOX concentrations; black circles are the average YFP and mScarlet values at each given DOX concentration (two replicates).

Further refuting the model, the rheostat-only *gal80Δ* strain shows a wide range of glucose-dependent chromatin accessibility (Fig. 2A, bottom right), despite always having an ON fraction close to 1, whereas the switch-only *mig1Δ* strain’s chromatin is either fully-open or fully-closed (Fig. 2A, top right), suggesting that not only the switch but also the Mig1p-mediated rheostat affects chromatin accessibility. We created a doxycycline titratable Mig1p strain to test this hypothesis (*mig1Δgal80Δ* TetO7pr-MIG1 *GAL1*pr-YFP; SLYM03 in Table S1, also Methods). Increasing Mig1p expression by increasing doxycycline lowered both the *GAL1*pr-YFP ON level (Fig. S3L) and the chromatin accessibility in a sugar-independent manner. Remarkably, the relationship between chromatin accessibility and transcription was the same as in a wild-type strain (Fig. 2A). Together, these data support an alternate mechanism for regulation of the decoupling and refute the hypothesis that only the switch acts on chromatin (Fig. S3A).

Our observation that chromatin accessibility correlates with ON level in a Mig1p-dependent fashion conflicts with previous reports that nucleosome and Mig1p binding are mutually exclusive at the *GAL1*pr (*9*). To investigate this apparent paradox, we deleted the Mig1p binding sites in the *GAL1*pr-YFP reporter (Fig. S4A) and measured the response to a glucose-galactose double gradient. Surprisingly, the phenotype of the *GAL1*pr^*mig1bsΔ*^ reporter strain was nearly identical to that of wildtype (Fig. 3B,C), despite the fact that the Mig1p binding sites are highly conserved and therefore expected to be functionally important (*12*). This indicates that the direct effect of Mig1p at *GAL1* is a relatively minor contribution at steady-state. Based on this, we concluded that the Mig1p rheostat must regulate *GAL1* upstream of the *GAL1* promoter.

**Fig. 3.**
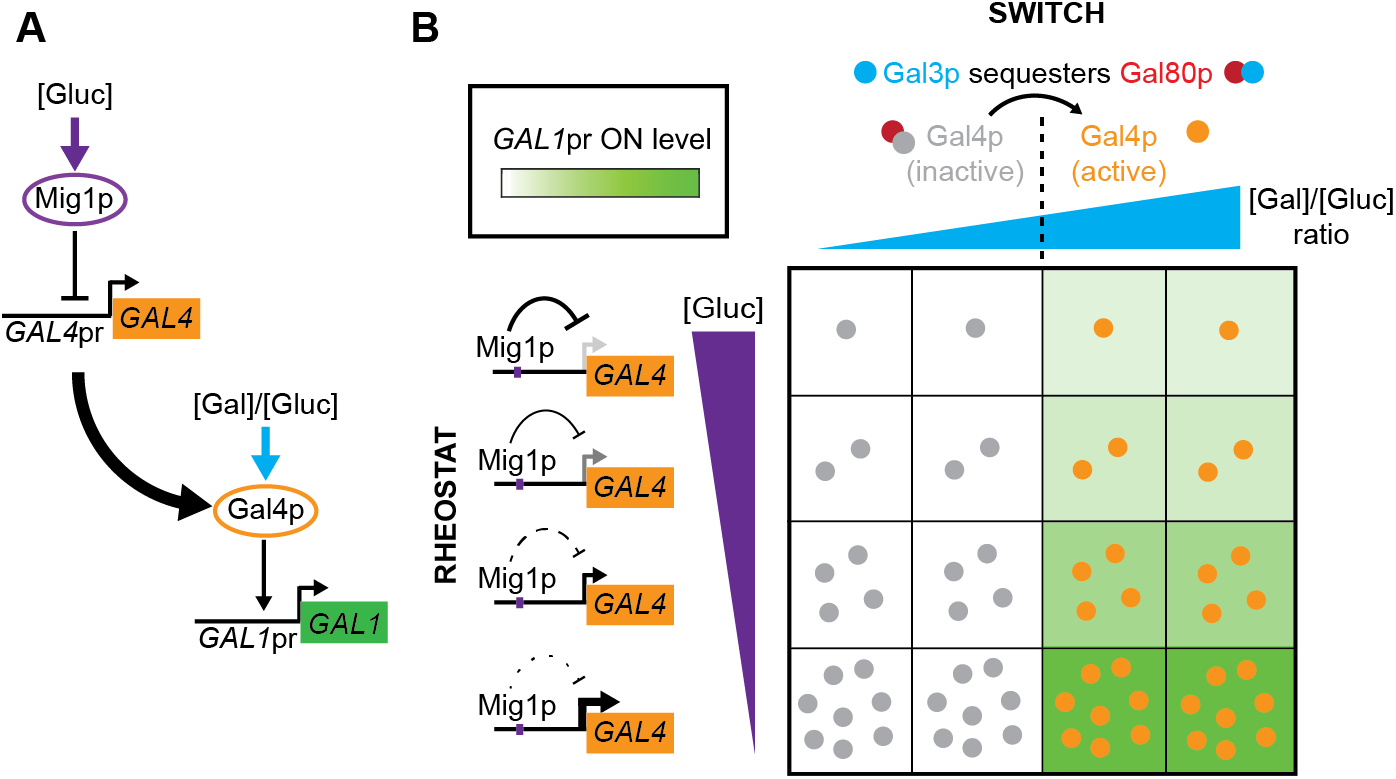
Mechanistic model: switch-and-rheostat works through decoupled regulation of Gal4p activity and level. (A) Based on our data, we propose that the switch-and-rheostat regulation in the GAL pathway is achieved through a hierarchical design; in response to glucose, the rheostat works via Mig1p transcriptional regulation of *GAL4,* where the activity of Gal4p is then regulated in response to the galactose:glucose ratio. (B) Molecular mechanism of the independent control of Gal4p level and activity. Internal galactose concentrations (which depend on the external ratio of glucose to galactose) controls the amount of activity of Gal3p (x-axis). Gal80p sequesters Gal4p; active Gal3p sequesters Gal80p. Thus, when the active Gal3p exceeds the total Gal80p concentration, Gal4p will convert sharply, in the galactose:glucose ratio space, from inactive (grey circles) to active (orange circles). But, the total amount of active Gal4p controls the amount of transcriptional output from the *GAL1* promoter (green shading). Glucose controls Mig1p activity thereby setting the total level of Gal4p (y-axis). Thus, glucose controls the level of Gal1p by controlling the total level of Gal4p and thereby the amount of active Gal4p.

Having ruled out Mig1p at the *GAL1* promoter as the mechanism of ON level control, we sought to determine the relevant Mig1p target. In addition to regulating *GAL1* expression, Mig1p is known to inhibit *GAL3* and *GAL4* expression (*6, 13*) (Fig. 1A), and there is evidence that changing Gal3p and Gal4p levels can affect the GAL response (*14, 15*). Surprisingly, deleting the Mig1p binding site in the *GAL4* promoter (Fig. S4B) was sufficient to phenocopy the behavior of a *mig1Δ* strain for steady-state *GAL1* expression (Fig. 3B,C), suggesting that glucose control of Gal1p levels is achieved solely through regulation of Gal4p levels. Supporting this hypothesis, YFP reporters for other Gal4p-regulated promoters, including two synthetic promoters, show similar glucose-mediated ON-level control even if they do not contain a Mig1p binding site (Fig. S4C-H).

To directly measure whether Gal4p abundance drives different *GAL1* expression levels and thus mediates the Mig1p rheostat, we built a strain with a doxycycline titratable mScarlet-I-Gal4p fusion (*16*) (*gal4Δ* TetO7pr-mScarlet-I-*GAL4*; SLYM08 in Table S1, Methods) (Fig. S4O,P). When we measured fluorescence from both mScarlet-I-Gal4p and *GAL1*pr-YFP, we observed a direct correlation between RFP and YFP levels in the ON subpopulation (Fig. 2D, Fig. S4K-N). As predicted, GAL pathway induction was highly sensitive to Gal4p level (see also Fig. S4I,J).

Our results support a mechanistic model where decoupling is achieved through the independent regulation of the level and activity of a single transcription factor, Gal4p (Fig. 3). Specifically, transcriptional regulation of Gal4p levels by Mig1p mediates the response to glucose, while protein binding of Gal80p to Gal4p (*17, 18*) regulates Gal4p activity in response to the galactose:glucose ratio. This “hierarchically-decoupled regulation” model can be contrasted with the “chromatin-decoupled regulation” model we hypothesized initially (Fig. S3A) in that only a single transcription factor, Gal4p, controls both switch and rheostat at the final output of the pathway. In both models decoupling is achieved via regulation working at two different levels, reminiscent of other cases in transcription regulation such as frequency versus amplitude modulation of the Msn2p/Msn4p stress responses in yeast (*19*).

We next investigated potential physiological advantages of decoupling the ON fraction from the ON level. When faced with mixtures of sugars, yeast first utilize glucose, then the less preferred carbon (*20*). This leads to a two-phase or diauxic growth (*21*). Yeast prepare by inducing GAL genes before glucose is depleted, the earlier a strain expresses GAL genes, the higher the fitness advantage it has once glucose is depleted. However, preparation comes at fitness cost before the glucose runs out (*22*). We hypothesized that one possible function of the decoupled switch-and-rheostat design is to lower the cost of preparation.

The rheostat lowers the cost of preparation. We competed a wild-type and rheostat-lacking *GAL4*pr^*mig1bsΔ*^ strain during diauxic-growth conditions. *GAL1*pr-YFP quickly reaches its maximum level in *GAL4*pr^*mig1bsΔ*^ strain while the level in a wildtype strain is still strongly dimmed until glucose is depleted (Fig. S5B,D). While glucose is being depleted, the wildtype strain has a clear fitness advantage over the GAL4pr^*mig1bsΔ*^ strain (Fig. 4A, Methods). This suggests that the excess cost of maximally inducing the GAL pathway (*22, 23*) in a *GAL4*pr^*mig1bsΔ*^ is not compensated for by any potential benefit of utilizing galactose. As glucose levels deplete below detection limit (Fig. S5B), wild-type GAL gene induction increases (Fig. S5D) and the *GAL4*pr^*mig1bsΔ*^ strain has a competitive advantage (t=7-10hr) (Fig. 4A). This advantage is temporary, persisting only until the unprepared strain induces a sufficient amount of galactose-responsive genes to take advantage of the galactose (*22*). This kinetic benefit was insufficient to overcome the initial steady-state fitness cost (Fig. 4A). The observed fitness differences are not due to difference in growth on glucose or galactose alone (Fig. 4A, Fig. S5F,G). Additionally, when cells are switched directly to galactose, wild type does not have a fitness advantage (Fig. 4A, Fig. S5E). In total, this evidence shows the rheostat reduces the fitness cost that is incurred by preparing for galactose utilization before glucose depletion.

**Fig. 4.**
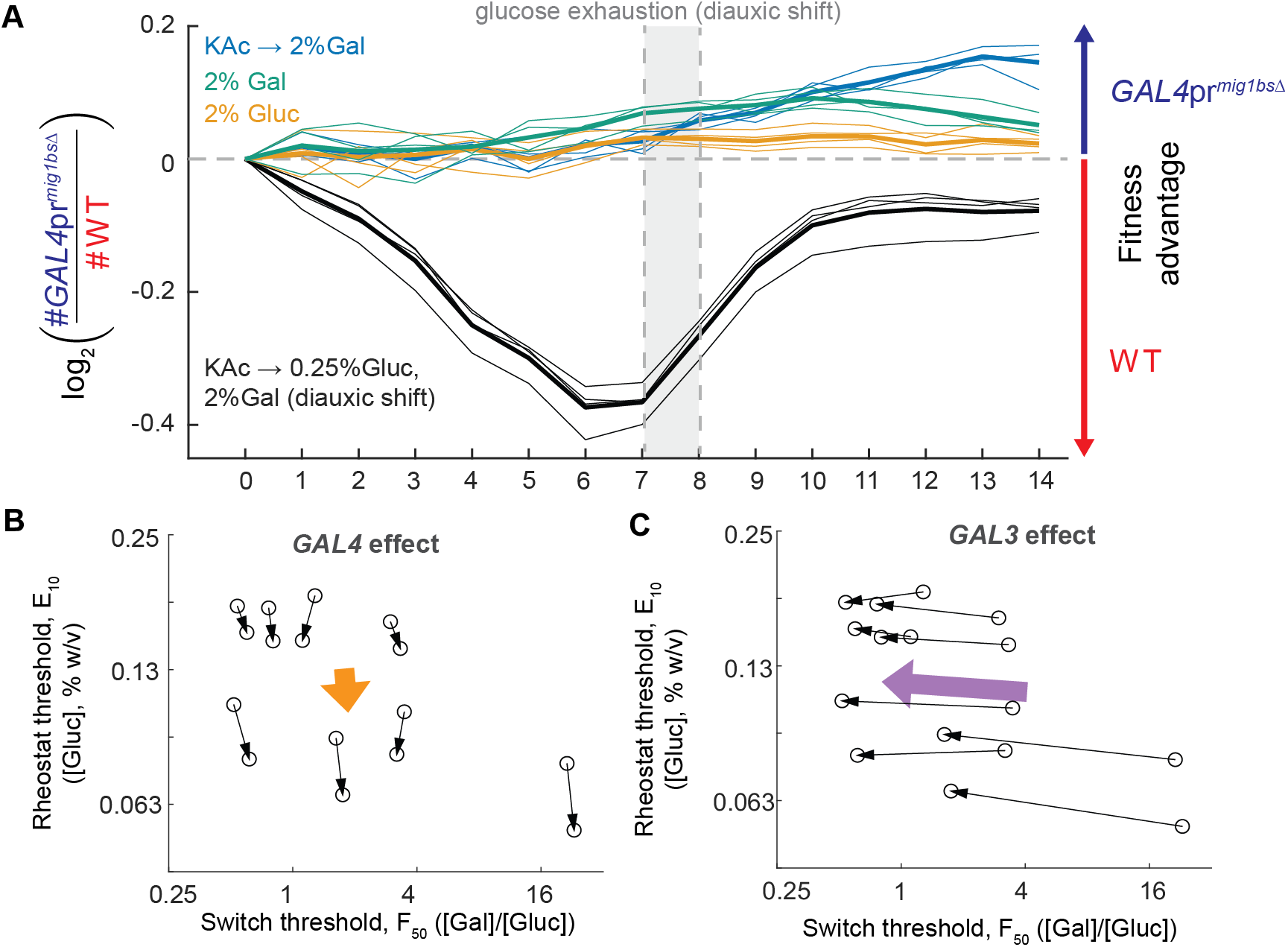
Physiological benefit and tunability through natural variation of switch-and-rheostat design. (A) Log_2_-ratio of *GAL4*pr^*mig1bsΔ*^ cell counts over wildtype cell counts versus time during either a diauxic growth (black, four replicate co-cultures), an “instantaneous” shift to pure galactose (blue, four replicate co-cultures), steady-state growth on 2% glucose (yellow, four replicate co-cultures), or steady-state growth on 2% galactose (green, four replicate co-cultures). Thick lines are the average (thick line) of the four measurements (thin lines). A positive value on the y-axis shows that *GAL4*pr^*mig1bs*Δ^ has a net fitness advantage since the initial timepoint, while a negative value indicates the wildtype has a net fitness advantage since the initial timepoint. (B)-(C) Each point represents the (F50,E10) thresholds of a strain with a specific combination of S288C or DBVP1106 alleles of *GAL3, GAL4*, and *MKT1* in either the S288C or DBVPG1106 background. Black arrows connect strains that only differ in their *GAL3* (C) or *GAL4* (D) allele. The purple and orange arrows represent the average effect of all *GAL3* and *GAL4* allele swaps respectively. *GAL3, GAL4,* and *MKT1* were identified as causative loci from bulk segregant analysis (Fig. S6).

We next sought to determine if natural isolates have genetic variants that independently affect the rheostat and the switch. We have shown that there is natural variation in both the rheostats and switch (*24, 25*). We chose two isolates, S288C and DBVPG1106 (Fig. S6A-C), and performed bulk-segregant linkage mapping using a pooled sorting strategy (Supplementary Text, Fig. S6D-G) to identify three loci that affect either the switch or the rheostat. Each locus had a candidate gene: *GAL3, MKT1*, and *GAL4* (Fig. S6G). We used CRISPR/cas9 to construct all combinations of strain backgrounds and alleles and measured the resulting phenotypic landscape (Fig. S6H). We found that *GAL4* affects the rheostat but not the switch, and vice versa for *GAL3* (Fig. 4B,C, Fig. S6K). Strain background and *MKT1* affect both (Fig. S6I-K). Thus, the switch and rheostat can be tuned independently of each other, highlighting the functional separability of induction decision and level.

Decoupled control is a useful property as it allows response features to be independently controlled physiologically and evolutionarily (*26–28*). While mechanistic dissection of decoupling is limited, it is often assumed to involve independent transcription factor binding sites on promoters and can be aided by chromatin (*8*). The mechanism we uncover here, of hierarchically-decoupled regulation of a transcription factor, is a ubiquitous motif (*29, 30*) and thus decoupling of response is likely ubiquitous in other systems. One advantage of decoupling achieved through this hierarchically-decoupled regulation is that all genes regulated by the subordinate transcriptional regulator will gain the regulation of the superior transcriptional regulator, allowing the decoupled response to be coordinated among all the genes. This design has the potential to make the evolution of complex coordinated regulation easier. Our results, combined with previous work, may provide guidance to what types of control features to look for given the biological behaviors that are expected. We encourage the community to examine whether these rules hold in other eukaryotic signaling systems.

## Acknowledgments

We thank Rebecca Ward, Dan Davidi, Ron Milo, and Allon Klein for critical feedback on the manuscript; Sarah Boswell, Severin Schink, Han-Ying Jhuang, Nathan Johnson, and members of the Springer Lab for helpful discussions; the HMS Systems Biology FACS Facility for technical support; and Shervin Javadi and Stratedigm for flow cytometry assistance.

## Author contributions

C.R.-T., J.W., I.B.-Z., J.P., A.L., Y.S., and M.S. designed the experiments. C.R.-T., J.W., I.B.-Z., J.P., and A.L. performed experiments and analyzed results. C.R.-T., J.W., I.B.-Z., and M.S. wrote the manuscript. All authors read and approved the final manuscript.

## Data and materials availability

All datasets generated and analyzed during the current study are available for download from Dryad. All custom code used during the current study is available for download from Dryad and Github (https://github.com/springerlab/Flow-Cytometry-Toolkit).

## Supplementary Materials

Materials and Methods

Supplementary Text

Figures S1-S7

Tables S1-S4

References (*31–47*)

## References and Notes

1. R. Escalante-Chong, Y. Savir, S. M. Carroll, J. B. Ingraham, J. Wang, C. J. Marx, M. Springer, Galactose metabolic genes in yeast respond to a ratio of galactose and glucose. Proc. Natl. Acad. Sci. U. S. A. 112, 1636–1641 (2015).

2. O. S. Venturelli, H. El-Samad, R. M. Murray, Synergistic dual positive feedback loops established by molecular sequestration generate robust bimodal response. Proc. Natl. Acad. Sci. U. S. A. 109, E3324–33 (2012).

3. P. J. Bhat, Galactose Regulon of Yeast: From Genetics to Systems Biology (Springer Science & Business Media, 2008).

4. V. Pilauri, M. Bewley, C. Diep, J. Hopper, Gal80 dimerization and the yeast GAL gene switch. Genetics. 169, 1903–1914 (2005).

5. A. Traven, B. Jelicic, M. Sopta, Yeast Gal4: a transcriptional paradigm revisited. EMBO Rep. 7, 496–499 (2006).

6. M. Johnston, J. S. Flick, T. Pexton, Multiple mechanisms provide rapid and stringent glucose repression of GAL gene expression in Saccharomyces cerevisiae. Mol. Cell. Biol. 14, 3834–3841 (1994).

7. K. S. Zaret, J. S. Carroll, Pioneer transcription factors: establishing competence for gene expression. Genes Dev. 25, 2227–2241 (2011).

8. F. H. Lam, D. J. Steger, E. K. O’Shea, Chromatin decouples promoter threshold from dynamic range. Nature. 453, 246–250 (2008).

9. E. Frolova, M. Johnston, J. Majors, Binding of the glucose-dependent Mig1p repressor to the GAL1 and GAL4 promoters in vivo: regulationby glucose and chromatin structure. Nucleic Acids Res. 27, 1350–1358 (1999).

10. J. D. Buenrostro, B. Wu, H. Y. Chang, W. J. Greenleaf, ATAC-seq: A Method for Assaying Chromatin Accessibility Genome-Wide. Curr. Protoc. Mol. Biol. 109, 21.29.1–9 (2015).

11. A. N. Schep, J. D. Buenrostro, S. K. Denny, K. Schwartz, G. Sherlock, W. J. Greenleaf, Structured nucleosome fingerprints enable high-resolution mapping of chromatin architecture within regulatory regions. Genome Res. 25, 1757–1770 (2015).

12. M. Kellis, N. Patterson, M. Endrizzi, B. Birren, E. S. Lander, Sequencing and comparison of yeast species to identify genes and regulatory elements. Nature. 423, 241–254 (2003).

13. P. A. Silver, L. P. Keegan, M. Ptashne, Amino terminus of the yeast GAL4 gene product is sufficient for nuclear localization. Proc. Natl. Acad. Sci. U. S. A. 81, 5951–5955 (1984).

14. D. W. Griggs, M. Johnston, Regulated expression of the GAL4 activator gene in yeast provides a sensitive genetic switch for glucose repression. Proc. Natl. Acad. Sci. U. S. A. 88, 8597–8601 (1991).

15. M. Acar, B. F. Pando, F. H. Arnold, M. B. Elowitz, A. van Oudenaarden, A general mechanism for network-dosage compensation in gene circuits. Science. 329, 1656–1660 (2010).

16. D. S. Bindels, L. Haarbosch, L. van Weeren, M. Postma, K. E. Wiese, M. Mastop, S. Aumonier, G. Gotthard, A. Royant, M. A. Hink, T. W. J. Gadella Jr, mScarlet: a bright monomeric red fluorescent protein for cellular imaging. Nat. Methods. 14, 53–56 (2017).

17. O. Egriboz, S. Goswami, X. Tao, K. Dotts, C. Schaeffer, V. Pilauri, J. E. Hopper, Self-association of the Gal4 inhibitor protein Gal80 is impaired by Gal3: evidence for a new mechanism in the GAL gene switch. Mol. Cell. Biol. 33, 3667–3674 (2013).

18. F. Jiang, B. R. Frey, M. L. Evans, J. C. Friel, J. E. Hopper, Gene activation by dissociation of an inhibitor from a transcriptional activation domain. Mol. Cell. Biol. 29, 5604–5610 (2009).

19. Z. AkhavanAghdam, J. Sinha, O. P. Tabbaa, N. Hao, Dynamic control of gene regulatory logic by seemingly redundant transcription factors. Elife. 5, 684 (2016).

20. J. M. Gancedo, Yeast carbon catabolite repression. Microbiol. Mol. Biol. Rev. 62, 334–361 (1998).

21. L. Galdieri, S. Mehrotra, S. Yu, A. Vancura, Transcriptional regulation in yeast during diauxic shift and stationary phase. OMICS. 14, 629–638 (2010).

22. J. Wang, E. Atolia, B. Hua, Y. Savir, R. Escalante-Chong, M. Springer, Natural variation in preparation for nutrient depletion reveals a cost-benefit tradeoff. PLoS Biol. 13, e1002041 (2015).

23. G. I. Lang, A. W. Murray, D. Botstein, The cost of gene expression underlies a fitness trade-off in yeast. Proc. Natl. Acad. Sci. U. S. A. 106, 5755–5760 (2009).

24. K. B. Lee, J. Wang, J. Palme, R. Escalante-Chong, B. Hua, M. Springer, Polymorphisms in the yeast galactose sensor underlie a natural continuum of nutrient-decision phenotypes. PLoS Genet. 13, e1006766 (2017).

25. J. Wang, J. Palme, K. B. Lee, M. Springer, Natural Genetic Variation Can Independently Tune the Induced Fraction and Induction Level of a Bimodal Signaling Response. bioRxiv (2017), p. 131938.

26. H. Mengistu, J. Huizinga, J.-B. Mouret, J. Clune, The Evolutionary Origins of Hierarchy. PLoS Comput. Biol. 12, e1004829 (2016).

27. H. Yu, M. Gerstein, Genomic analysis of the hierarchical structure of regulatory networks. Proc. Natl. Acad. Sci. U. S. A. 103, 14724–14731 (2006).

28. D. M. Lorenz, A. Jeng, M. W. Deem, The emergence of modularity in biological systems. Phys. Life Rev. 8, 129–160 (2011).

29. S. S. Shen-Orr, R. Milo, S. Mangan, U. Alon, Network motifs in the transcriptional regulation network of Escherichia coli. Nat. Genet. 31, 64–68 (2002).

30. R. Milo, S. Shen-Orr, S. Itzkovitz, N. Kashtan, D. Chklovskii, U. Alon, Network motifs: simple building blocks of complex networks. Science. 298, 824–827 (2002).

31. R. Daniel Gietz, R. A. Woods, in Methods in Enzymology, C. Guthrie, G. R. Fink, Eds. (Academic Press, 2002), vol. 350, pp. 87–96.

32. C. Janke, M. M. Magiera, N. Rathfelder, C. Taxis, S. Reber, H. Maekawa, A. Moreno-Borchart, G. Doenges, E. Schwob, E. Schiebel, M. Knop, A versatile toolbox for PCR-based tagging of yeast genes: new fluorescent proteins, more markers and promoter substitution cassettes. Yeast. 21, 947–962 (2004).

33. W. M. Shaw, H. Yamauchi, J. Mead, G.-O. F. Gowers, D. J. Bell, D. Öling, N. Larsson, M. Wigglesworth, G. Ladds, T. Ellis, Engineering a Model Cell for Rational Tuning of GPCR Signaling. Cell. 177, 782–796.e27 (2019).

34. M. E. Lee, W. C. DeLoache, B. Cervantes, J. E. Dueber, A Highly Characterized Yeast Toolkit for Modular, Multipart Assembly. ACS Synth. Biol. 4, 975–986 (2015).

35. M. Baym, S. Kryazhimskiy, T. D. Lieberman, H. Chung, M. M. Desai, R. Kishony, Inexpensive multiplexed library preparation for megabase-sized genomes. PLoS One. 10, e0128036 (2015).

36. X. Chen, Y. Shen, W. Draper, J. D. Buenrostro, U. Litzenburger, S. W. Cho, A. T. Satpathy, A. C. Carter, R. P. Ghosh, A. East-Seletsky, J. A. Doudna, W. J. Greenleaf, J. T. Liphardt, H. Y. Chang, ATAC-see reveals the accessible genome by transposase-mediated imaging and sequencing. Nat. Methods. 13, 1013–1020 (2016).

37. L. Keren, O. Zackay, M. Lotan-Pompan, U. Barenholz, E. Dekel, V. Sasson, G. Aidelberg, A. Bren, D. Zeevi, A. Weinberger, U. Alon, R. Milo, E. Segal, Promoters maintain their relative activity levels under different growth conditions. Mol. Syst. Biol. 9, 701 (2013).

38. T. Hashida-Okado, A. Ogawa, I. Kato, K. Takesako, Transformation system for prototrophic industrial yeasts using the AUR1 gene as a dominant selection marker. FEBS Lett. 425, 117–122 (1998).

39. S. Treusch, F. W. Albert, J. S. Bloom, I. E. Kotenko, L. Kruglyak, Genetic mapping of MAPK-mediated complex traits Across S. cerevisiae. PLoS Genet. 11, e1004913 (2015).

40. F. A. Cubillos, L. Parts, F. Salinas, A. Bergström, E. Scovacricchi, A. Zia, C. J. R. Illingworth, V. Mustonen, S. Ibstedt, J. Warringer, E. J. Louis, R. Durbin, G. Liti, High-resolution mapping of complex traits with a four-parent advanced intercross yeast population. Genetics. 195, 1141–1155 (2013).

41. M. D. Edwards, D. K. Gifford, High-resolution genetic mapping with pooled sequencing. BMC Bioinformatics. 13 Suppl 6, S8 (2012).

42. F. W. Albert, S. Treusch, A. H. Shockley, J. S. Bloom, L. Kruglyak, Genetics of single-cell protein abundance variation in large yeast populations. Nature. 506, 494–497 (2014).

43. J. E. DiCarlo, J. E. Norville, P. Mali, X. Rios, J. Aach, G. M. Church, Genome engineering in Saccharomyces cerevisiae using CRISPR-Cas systems. Nucleic Acids Res. 41, 4336–4343 (2013).

44. A. A. Horwitz, J. M. Walter, M. G. Schubert, S. H. Kung, K. Hawkins, D. M. Platt, A. D. Hernday, T. Mahatdejkul-Meadows, W. Szeto, S. S. Chandran, J. D. Newman, Efficient Multiplexed Integration of Synergistic Alleles and Metabolic Pathways in Yeasts via CRISPR-Cas. Cell Syst. 1, 88–96 (2015).

45. A. M. Deutschbauer, R. W. Davis, Quantitative trait loci mapped to single-nucleotide resolution in yeast. Nat. Genet. 37, 1333–1340 (2005).

46. S.-I. Lee, A. M. Dudley, D. Drubin, P. A. Silver, N. J. Krogan, D. Pe’er, D. Koller, Learning a prior on regulatory potential from eQTL data. PLoS Genet. 5, e1000358 (2009).

47. H. Sinha, B. P. Nicholson, L. M. Steinmetz, J. H. McCusker, Complex genetic interactions in a quantitative trait locus. PLoS Genet. 2, e13 (2006).

