## Supplementary Materials for "Decoupling transcription factor expression and activity enables dimmer-switch gene regulation"

**This PDF file includes:**

Materials and Methods
Supplementary Text
Figs. S1 to S7
Table S4

**Other Supplementary Materials for this manuscript include the following:**

Tables S1-S3

### Materials and Methods

Strains and Reporter Construction

All strains used in this study are listed in Table S1. The primary wild-type strain background used in this paper is haploid prototrophic S288C, specifically either FY4 (MATa) or FY5 (MATα) (*1*). The natural isolate strain DBVPG1106 was also used in this study(*22*). Deletions of *MIG1* and *GAL80* were performed by standard lithium acetate yeast transformation (*31*) with drug-maker deletion cassettes (*32*). Strain SLYM03 was built by transforming a prototrophic diploid (made from mating FY4 and FY5) with the *GAL1*pr-YFP reporter, MIG1 deletion cassette, and Tet-inducible MIG1 cassette, sporulating and selecting for MATa haploids containing all three constructs, mating to a MATα *gal80Δ* strain, then again sporulating and screening for a haploid strain containing all four constructs. In addition to traditional drug-marker-based strain construction, we also adapted a CRISPR/Cas9 genome editing strategy (*33*) to make markerless edits to the genome, e.g. disrupting the Mig1p binding sites in the *GAL4* promoter sequence (SLYM04). Plasmids for this purpose were constructed using the Yeast MoClo Toolkit (YTK) (*34*).

Growth Conditions

For all experimental assays, cells were grown in a mixture of synthetic minimal medium (S), which consists of 1.7g/L yeast nitrogen base and 5g/L ammonium sulfate, and a carbon source of choice. For incubations, cells were grown at 30ºC in an incubator with rotary shaking at 230 rpm for cultures in tubes/flasks, or a humidified incubator with rotary shaking at 999 rpm for cultures in 96-well 1.2-mL plates (Thermo Fisher 260251).

All strains were struck from glycerol stocks onto YPD agar and grown to form colonies, and then inoculated from single colonies into liquid YPD medium and grown overnight until saturation. These cultures were then used to inoculate a dilution series of outgrowth media (S+2% raffinose or S+2% glucose) and grown for 16-18hrs until reaching $OD_{600}$ 0.1-0.3.

Strains that are *gal80*Δ lack a source of GAL pathway repression and are constitutively induced when grown in raffinose. For this reason, glucose was used instead as the outgrowth sugar for *gal80*Δ strains to achieve an uninduced outgrowth condition. With this exception, raffinose was otherwise used as a non-repressive, non-inducing outgrowth sugar.

Glucose-galactose double gradient assay

To prepare a "double gradient" of galactose:glucose concentrations, glucose and galactose were serially diluted 2-fold across the rows and columns, respectively, of a 96-well 1.2mL plate in an S medium background. Selected outgrowth cultures were pelleted, washed twice with S medium (no carbon source), and then resuspended in S medium for inoculation 1:100 into a double gradient plate. This inoculum size has been previously shown to be low enough to not significantly affect the sugar concentrations during the experiment (*1*). These cultures were incubated for 8hr (or 24hrs with a back-dilution after 12hrs for *gal80*Δ strains to maintain the low culture density) to reach steady-state expression levels, then harvested by pelleting, washing twice, and resuspending in Tris-EDTA pH 7.5 (TE) before being run on the flow cytometer.

Glucose single gradient assay

For comparing the decision and level setpoints between strains, GAL induction was performed in a 2-fold dilution series of glucose concentration, from 1% to 0.002% w/v, with constant 0.25% galactose. To assess and control for well-to-well variation, these assays were performed as a co-culture of a "query" strain to be phenotyped and a "reference" strain that was always natural isolate YJM978 with constitutive mCherry segmentation marker (SLYB93).

Flow cytometry and data analysis

All fluorescence measurements were made using a Stratedigm S1000EX flow cytometer paired with an A600 autosampler and A700 plate-handling hotel.

FACS sorting was performed with a Sony SH800Z Cell Sorter using manual polygonal gates based on FSC v. FITC height values (Fig. S7). The sample and collection chambers of the sorter were kept at 4ºC, and all samples were held on ice before and after sorting.

Flow cytometry data were analyzed using custom MATLAB code (https://github.com/springerlab/Flow-Cytometry-Toolkit). As a general workflow, raw FCS data files were processed by manually gating out debris based on FSC/SSC measurements, normalizing fluorescence values to cell size by dividing by SSC values, and then performing expression analysis and other computations on the $\log_{10}$-transformed fluorescence data.

Peak calling algorithm using Gaussian fitting

To quantify the observation that along a glucose-galactose double gradient the fraction of induced cells is varied independently of the induction level of induced cells, we fitted the cell-size normalized logarithm of YFP expression histogram to one or two Gaussians. We first tried to fit a single Gaussian and only if goodness of fit was not high enough ($R^2 < R^2{}_{threshold}$) we fitted two Gaussians. We used an ON level threshold parameter ($ON_{threshold}$) to dictate if a unimodal population is ON or OFF and to restrict the Gaussian means in the fitting algorithm. Choice of fit parameters ($ON_{threshold}$, $R^2{}_{threshold}$) was done in a way that minimized discrepancy from manual calling (See Fig. S2M for Gaussian fitting without restricting the fit parameters).

For most of the analysis in this work, we chose the described Gaussian fitting metric, and not the area metric we used previously, because this new method identifies ON and OFF subpopulations within bimodal populations without assuming anything about the OFF population. The area metric which we previously used (*1*) to calculate the fraction of induced cells relied on subtracting the OFF histogram, defined as the histogram of each strain in 1% glucose. This method led us to previously overlook the fact that all *gal80Δ* samples are ON compared to the wildtype in 1% glucose. A correction would be to define the OFF histogram once for the wildtype, but this would cause day to day differences in the flow cytometer readings to strongly affect the conclusions. Our current method is similar to the one used by Venturelli et al. (*2*) to distinguish bimodal from unimodal samples. See comparison between the three metrics in Fig. S2K-N.

ATAC-seq assay and promoter capture

We adapted the protocol of Schep et al. (*11*) to perform ATAC-seq on crosslinked yeast samples. At the point of harvesting, samples were crosslinked for 30min with 1% formaldehyde, quenched for 10min with the addition of 2.5M glycine, then pelleted and washed with ice-cold PBS. If sorted on the same day, pelleted cells were held on ice until sorted; otherwise, these samples were snap-frozen in a dry ice-ethanol bath and stored at -80ºC until ready to sort. Sorted samples were likewise stored at -80ºC until ready to be processed in parallel.

To begin ATAC-seq library preparation, all samples (10^6 sorted cells, or the equivalent OD for non-sorted samples) were reformatted into DNA lo-bind 96-well PCR plates. For lysis, samples were incubated with 0.25U/ul Zymolyase 100T for 15min at 30C, then spun for 5min at 6100xg, 4ºC to pellet cells/genomic DNA while discarding the supernatant. Using Nextera reagents (Illumina FC-121-1031), samples were then tagmented with 10-fold diluted Tn5 enzyme in TD buffer for 15min at 37ºC, modified from a protocol for low-input library preparation(*35*). To stop the tagmentation reaction and reverse the crosslinking, a concentrated "reverse crosslink solution" was added to final concentrations of 1%SDS, 1mM EDTA, 0.33mg/mL proteinase K, and 0.2M NaCl, modified from Chen et al.(*36*), and incubated overnight for 12-16hr at 65ºC. DNA was then purified using 1.6X SPRI beads, PCR amplified for 12 cycles using customized Nextera indices from Baym et al. (*35*), and pooled equimolarly based on quant-iT (Thermo Fisher Q33120) measurements to form the pre-capture ATAC-seq library.

In order to increase sequencing depth at regions of interest, we selected 17 genes (both control and experimental) for promoter capture: *ACT1*, *CAT8*, *GAL1*, *GAL2*, *GAL3*, *GAL4*, *GAL7*, *GAL80*, *GCY1*, *HXT1*, *MAL31*, *MAL33*, *SUC2*, *TUB1*, *RPS14B*, *EFB1*, and *GAS1*. To enrich for these promoters of interest, a pool of biotinylated DNA probes was created by PCR amplifying the targeted regions (Table S3) with biotin-dCTP and shearing the resulting PCR products to a size range of 150-300bp using a Covaris S2 sonicator, as verified by running on an Agilent 2100 Bioanalyzer. The library was hybridized to the DNA capture probes, pulled down using M270 streptavidin beads, amplified for 12 cycles using Illumina primers, and purified using SPRI beads to form a post-capture ATAC-seq library. The library was sequenced on an Illumina NextSeq 500 sequencer. As a result of the capture, we observed fold enrichment of approximately 50-100x (Fig. S3C) in coverage at regions of interest relative to the rest of the genome, allowing for the pooled sequencing of 100s of samples in a single sequencing run.

Analysis of ATAC-seq data

Alignments were mapped to the S288C reference genome assembly R64 (sacCer3) using bowtie2 and sorted using samtools. Any reads with identical start and stop positions, presumably PCR duplicates generated during the library preparation, were computationally discarded to minimize PCR amplification biases that might result from the additional amplification steps following hybrid capture. To normalize for input size of each sample into the library, the summed number of reads at control loci *TUB1*, *ACT1*, *GAS1*, *EFB1*, and *RPS14B*, selected based on their invariant expression over different growth conditions (*37*), was used to normalize the start site counts at the *GAL1* promoter. To compare between strain backgrounds, all summed counts were normalized by subtraction to those of wildtype cells grown in 2%glucose (defined as 0% chromatin accessibility) and divided by the difference in summed counts between wildtype cells grown in 2% galactose (defined as 100% chromatin accessibility) and those in 2% glucose.

Microscopy

Fluorescence microscopy was performed using a Nikon Ti-E microscope with Prior Lumen 200 Pro light source, Hamamatsu ORCA-R2 CCD camera and Plan Apo VC 60x/1.40 objective. Image segmentation and quantitation was performed using custom Python scripts, with local otstu thresholding based on YFP signal used to define boundaries of individual cells and, for each cell region defined in this way, calculating the average YFP and RFP intensity per pixel.

Diauxic fitness competition measurement

As in Wang et al. (*22*), strains were grown in co-culture beginning from the inoculation of the outgrowth cultures (potassium acetate, glucose or galactose), through to the final inoculation of the experimental cultures (glucose+galactose, glucose or galactose). At hourly timepoints, aliquots of culture were taken off for measuring via flow cytometry and fixed by the addition of sodium azide to a final concentration of 0.1% azide in TE. Additionally, media was collected at each timepoint by spinning down aliquots of culture and taking off the cell-free supernatant. Sugar measurements then were made by adding a sugar-specific oxidase (Megazyme K-GLUHK-110A, K-ARGA) and measuring absorbance at 340nm as in Wang et al. (*22*).

Crossing and generating segregants

To prepare parent strains for crossing and sporulation, we sporulated diploid natural isolates bearing the ho::*GAL1*pr-YFP-hphNT1 reporter cassette and isolated random spores that displayed MATa or MATα phenotypes in test crosses. We then introduced a constitutive fluorescent marker in tandem with the GAL reporter, to obtain MATa; ho::*GAL1*pr-YFP-mTagBFP2-kanMX4 or MATα; ho::*GAL1*pr-YFP-mCherry-natMX4 parent strains. To the MATa parent we also introduced a pRS413-derived plasmid bearing STE2pr-AUR1-C and hphNT1. This plasmid is maintained by hygromycin selection but also allows selection for MATa cells by Aureobasidin A (*38*). This plasmid design is inspired by a similar mating-type selection plasmid used in a recent study (*39*).

To isolate segregants for phenotyping, we crossed a parent with BFP-kanMX + MAT-selection plasmid to a parent with mCherry-natMX and isolated a G418RNatRHygR diploid hybrid with the plasmid. We sporulated the hybrid by culturing it to saturation in YPD, diluting 1:10 in YP+2% potassium acetate and incubating at 30C for 8 hours, and washing and resuspending into 2% potassium acetate and incubating at 30C until >20% of cells were tetrads, or about 3 days. We incubated ~$5x10^6$ tetrads in 100uL water with 50U of zymolyase 100T (Zymo Research) for 5 hours at 30C, and then resuspended tetrads in 1mL of 1.5% NP-40 and sonicated for 10 seconds at power setting 3 on a probe sonicator (Fisher Scientific). The resulting segregants were plated on YPD + 0.5ug/mL Aureobasidin A ("AbA"; Clontech) and random colonies were picked into YPD liquid and saved as glycerol stocks. Haploidy was confirmed by mating to tester strains with known mating type. 90 segregants were phenotyped for GAL induction as described above.

Sorting-based bulk segregant analysis

To generate segregant pools, we prepared a diploid hybrid and sporulated it as described above. To reduce the size of recombination blocks and improve the resolution of linkage mapping (*40*), we then performed the following "intercross" protocol 4 times: from spore suspension, use Sony SH800 Cell Sorter to sort 4x106 BFP+ or mCherry+ (but not +/+ or -/-) cells into 100uL YPD + 40ug/mL tetracycline; incubate for 16 hours at 30C without shaking; add 5mL YPD + 200ug/mL G418 + 100ug/mL ClonNat + 200ug/mL Hygromycin B and incubate 48 hours at 30C with shaking; sporulate cultures and prepare sonicated spore suspension. After the 4th sporulation cycle, the sonicated spores were resuspended in YPD + 0.5ug/mL AbA and incubated at 30C for 16 hours. This culture was frozen as a glycerol stock, as well as used to inoculate the galactose-induction sorting experiment.

To sort segregant pools for bulk genotyping, we inoculated the intercrossed, MATa-selected segregants from a saturated YPD culture into S + 2% raffinose + AbA at dilutions of 1:200, 1:400, 1:800, and 1:1600, and incubated at 30C for 16-24 hours. We chose the raffinose culture with OD closest to 0.1, washed once in S (0.17% Yeast Nitrogen Base + 0.5% Ammonium Sulfate), and diluted 1:200 into S + 0.25% glucose + 0.25% galactose + AbA. We incubated the glucose-galactose culture at 30C for 8 hours, and then used a Sony SH800 sorter to isolate pools of 30,000 cells with the 5% lowest ("OFF") and highest ("HI") YFP expression, among cells whose Back Scatter (BSC) signal was between 105 and $3 \times 10^5$. The "LOW" pool was similarly obtained, but from the 5% of cells with lowest non-basal expression (Figure S7C). The sorted cells were resuspended in YPD + AbA and incubated at 30C until saturation, about 16-24 hours. An aliquot of this culture was saved for -80C glycerol stocks, and another was used to prepare sequencing libraries.

To sequence the segregant pools, we extracted genomic DNA from 0.5mL of saturated YPD culture of each segregant pool using the PureLink Pro 96 kit (Thermo Fisher K182104A). From these genomic preps, we made sequencing libraries using Nextera reagents (Illumina FC-121-1030) following a low-volume protocol (*35*). We adjusted the input DNA concentration so that resulting libraries had mean fragment sizes of 200-300bp as measured on a BioAnalyzer. Libraries were multiplexed and sequenced in an Illumina NextSeq flow cell to a depth of 16-33x.

Reads from the Illumina sequencing were aligned to the sacCer3 reference genome using bwa mem, and SNP counts were generated using samtools mpileup, on the Harvard Medical School O2 cluster. These outputs were processed in MATLAB using custom code as follows: SNPs with coverage less than 2 or more than 1000 were removed. The LOW and HI pools were computationally merged into an ON pool. To make sure the two pools contributed equally to the merged pool, at each SNP, allele counts in the pool with higher coverage were randomly subsampled to the coverage of the other pool. The final allele counts in each pool were output to text files by chromosome and given as inputs to the MULTIPOOL algorithm (mp_inference.py version 0.10.2) (*41*) to compute LOD scores. Loci with maximal LOD>5 were considered significant; previous work showed that this corresponded to an FDR of 5% (*39*, *42*). This correspondence may differ under our experimental conditions; therefore, the 2 loci that we did not validate experimentally should be interpreted with caution.

CRISPR/Cas9 allele replacement

Allele replacement strains were constructed using 3 rounds of gene knockout followed by CRISPR/Cas9-mediated markerless integration of heterologous allele. Initially, strains were prepared by introducing Cas9 on a CEN/ARS plasmid (SLVF11); this plasmid is derived from a previous one(*43*), but we replaced the auxotrophic *URA3* marker with AUR1-C to allow Aureobasidin A selection on prototrophic natural isolates. In each round of allele replacement, a gene plus upstream and downstream flanking sequences (-784bp to +815bp for *GAL3*, -449bp to +372bp for *MKT1*, -191bp to +139bp for *GAL4*) was deleted by integration of a kanMX6 marker with 40bp flanking homology. Then, a donor DNA, a guide RNA insert, and a guide RNA backbone were simultaneously transformed into the strain(*44*). The donor DNA contains the new allele, its flanking sequences, and an additional 40bp of homology to target it to the correct genomic locus. The guide RNA insert was a linear DNA fragment containing a SNR52 promoter driving a guide RNA gene containing a 20bp CRISPR/Cas recognition sequence linked to a sgRNA scaffold sequence, plus 40bp of flanking homology on both ends to a guide RNA backbone. The guide RNA backbone was a 2u plasmid containing natMX4 (pRS428). This was linearized by NotI + XhoI digestion before transformation. Allele re-integration transformations were plated on cloNAT to select for in vivo assembly of the guide RNA into a maintainable plasmid, and Aureobasidin A to select for presence of Cas9. Successful re-integration was verified by colony PCR and Sanger sequencing was performed on a subset of strains and on all donor DNAs to verify the sequence of allelic variants.

### Supplementary Text

Identifying causative alleles by bulk-segregant analysis

To identify these genes responsible for variation in $E_{10}$ and $F_{50}$ between strains S288C and DBVPG1106, we performed bulk-segregant linkage mapping using a pooled sorting strategy (Figure S6D). We grew a pooled mixture of haploid (MATa) segregants in a mixture of glucose and galactose where the parental *GAL1*pr-YFP distributions were maximally different (Figure S6D, grey). We then used FACS to sort the segregants into pools of uninduced ("OFF"), induced with low-expression ("LOW"), and induced with high-expression ("HIGH") cells (Figure S6D), and sequenced each pool. We expected that a genomic locus affecting ON level (and thus $E_{10}$) would differ in allele frequency between the LOW and HIGH pools, while any locus affecting ON fraction (and thus $F_{50}$) would differ in allele frequency between the OFF pool and a computationally-merged LOW+HIGH (ON) pool.

Three of the most significant loci, defined as genomic regions with a peak log-odds (LOD) score>5 calculated by MULTIPOOL (*41*), are centered at chrIV:457Kb, chrXIV:457Kb, and chrXVI:81Kb (Figure S6E). Scanning these loci, we found a single gene that seemed likely to be causative in each case. Two loci contained *GAL3* (chrIV) and *GAL4* (chrXVI) which are direct regulators of galactose sensing; the other contained *MKT1* (chrXIV) which is known to have pleiotropic effects in crosses between S288C and natural isolates (*45-47*). The *GAL3*-associated locus was significant only in the OFF/ON comparison, while the *GAL4*-associated locus was only significant in the LOW/HIGH comparison, suggesting that the effect of these loci are specific to $F_{50}$ and $E_{10}$ respectively. The *MKT1*-associated locus was significant in both comparisons but had a higher LOD score in the LOW/HIGH than in the OFF/ON comparison.

.

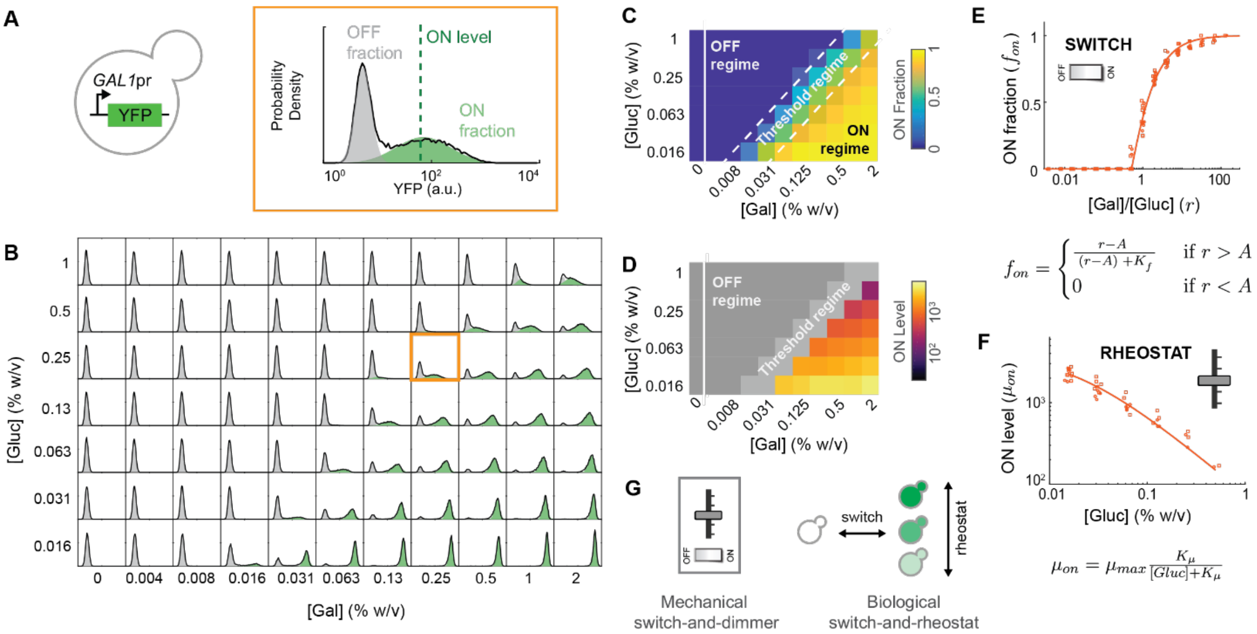


**Fig. S1.**

Response of the yeast GAL pathway to glucose-galactose combinations can be decomposed into a "switch" and "rheostat". (A) Diagram of *GAL1*pr-YFP reporter strain for GAL pathway activity and a representative histogram of the flow cytometry based measurement of the expression distribution. To extract metrics, the log(expression) distribution is fit to either one or two Gaussians (OFF and ON Gaussians respectively depicted in grey and green, Methods). The ON fraction is defined as the percent of the population in the ON Gaussian, and ON level is defined as the mean of the ON Gaussian. (B) The steady-state expression distribution of the *GAL1*pr-YFP reporter was measured across a double gradient of glucose (y-axis) and galactose (x-axis), histograms are of one of the two replicates (see Fig. S2A for overlay of both biological replicates measured). The orange square is the same histogram as in (A). (C) Heatmap of the ON fraction ($f_{ON}$) across the glucose-galactose double gradient, values are the average of two replicates for each glucose-galactose combination. We separated the response into three regimes: an "OFF regime", a "threshold regime", and an "ON regime" based on the ON fraction (Methods). (D) A plot of the ON fraction of all the experiments in (B) versus the galactose:glucose ratio is well fit by a shifted Michaelian function (circles indicate replicate 1, squares indicate replicate 2). X-values randomly jittered for visualization purposes. Fit shown uses $A = 0.52 \pm 0.08$ and $K_f = 0.77 \pm 0.08$, with fit s.e. = 0.04. (E) Heatmap of the ON level ($\mu_{ON}$) of cells from the ON regime in (C), values are the average of two replicates for each glucose-galactose combination. (F) A plot of ON level from the ON regime (E) versus glucose concentration is well fit by a Michaelian function (circles indicate replicate 1, squares indicate replicate 2). X-values randomly jittered for visualization purposes. Fit shown uses $\mu_{max} = 4{\cdot}10^3 \pm 1{\cdot}10^3$ and $K_\mu = 0.019 \pm 0.08$, with fit s.e. = 2·102. (G) Diagram of a mechanical switch-and-dimmer and our biological switch-and-rheostat.

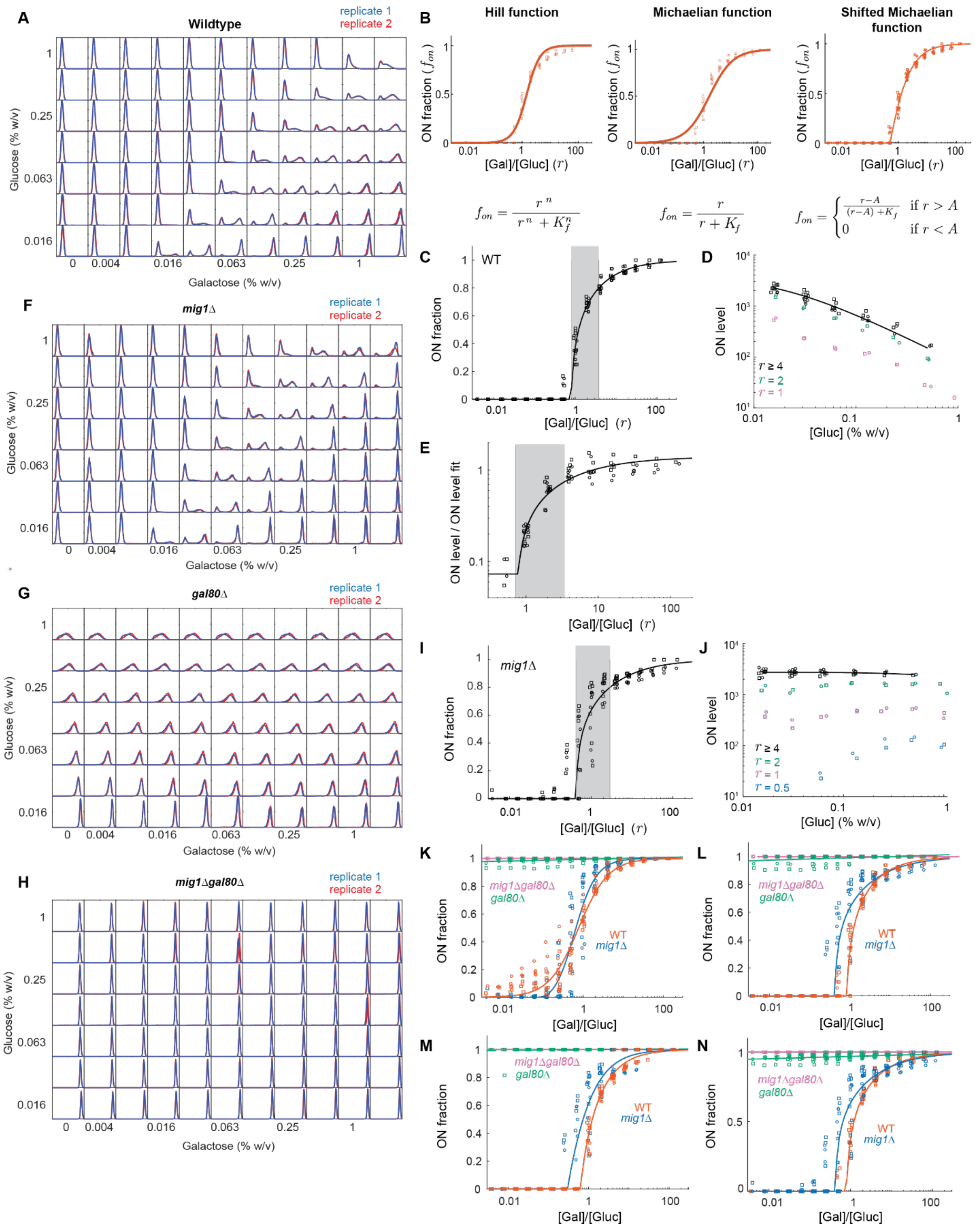

Wildtype
replicate 1
replicate 2
Glucose (% w/v)
Galactose (% w/v)
Hill function
Michaelian function
Shifted Michaelian function
ON fraction ($f_{on}$)
[Gal]/[Gluc] ($r$)
$f_{on} = \frac{r^n}{r^n + K_f^n}$
$f_{on} = \frac{r}{r + K_f}$
$f_{on} = \begin{cases} \frac{r-A}{(r-A)+K_f} & \text{if } r > A \\ 0 & \text{if } r < A \end{cases}$
WT
ON fraction
ON level
[Gluc] (% w/v)
ON level / ON level fit
mig1Δ
gal80Δ
mig1Δgal80Δ
[Gal]/[Gluc]


**Fig. S2.**

**Fig. S2. (Continued)**

Reproducibility, alternative fitting, and ON level dependence in the threshold regime for wildtype, *mig1*Δ, *gal80*Δ, and *mig1*Δ*gal80*Δ strains, related to Fig. 1 and Fig. S1. (A) Overlay of *GAL1*pr-YFP histograms along a glucose-galactose double gradient for the two biological repeats of the wildtype strain. For the rest of the figure squares summarize the data from replicate 1 and circles replicate 2. (B) ON fraction fitted by either a Hill function ($K_f = 1.45 \pm 0.07$, $n = 1.9 \pm 0.1$, fit s.e. = 0.06), a Michaelian function ($K_f = 1.6 \pm 0.2$, fit s.e. = 0.10), or a shifted Michaelian (reproduced for comparison from Fig. 1E), which has the best goodness of fit (fit s.e. = 0.04). (C) ON fraction versus galactose:glucose ratio ($r$) reproduced from Fig. 1D, with threshold regime highlighted in grey. (D) ON level versus glucose concentration (from the ON regime, $r \geq 4$) reproduced from Fig. S1F (black), overlaid with ON level from the threshold regime ($1 \leq r \leq 2$) (pink $r = 1$, green $r = 2$). (E) ON level in both threshold and ON regimes divided by ON level fit from the ON regime (from Fig. S1F). Its logarithm is well fit by the shifted Michaelian function from Fig. S1E, shifted and scaled to the relevant range. (F)-(H) Overlay of *GAL1*pr-YFP histograms along a glucose-galactose double gradient for the two biological repeats of the *mig1*Δ (F), *gal80*Δ (G) and *mig1*Δ*gal80*Δ (H) strains. (I) ON fraction versus galactose:glucose ratio ($r$) of a *mig1*Δ strain, reproduced from Fig. 1J, with the threshold regime highlighted in grey. (J) ON level versus glucose concentration (from the ON regime, $r \geq 4$) reproduced from Fig. 1K (black), overlaid with ON level from the threshold regime ($0.5 \leq r \leq 2$) (blue, pink, green). (K)-(N) Comparison between four metrics for calling the ON subpopulation, with resulting ON fraction values fit to a shifted Hill function for illustrative purposes. (K) Area metric from Escalante-Chong et al. (*1*). (L) Gaussian fitting starting from two Gaussians and moving on to a single Gaussian if the difference between the two means is less than 3 times the standard variation of the OFF peak, or if one of the weights is smaller than 0.05 (see Ref.2). (M) Gaussian fitting starting from a single Gaussian and moving to two gaussians if $R^2$ is smaller than 0.95, as we did for Fig. 1, but without restricting the fit parameters to the OFF and ON regions. (N) Gaussian fitting used in the main text for Fig. 1.

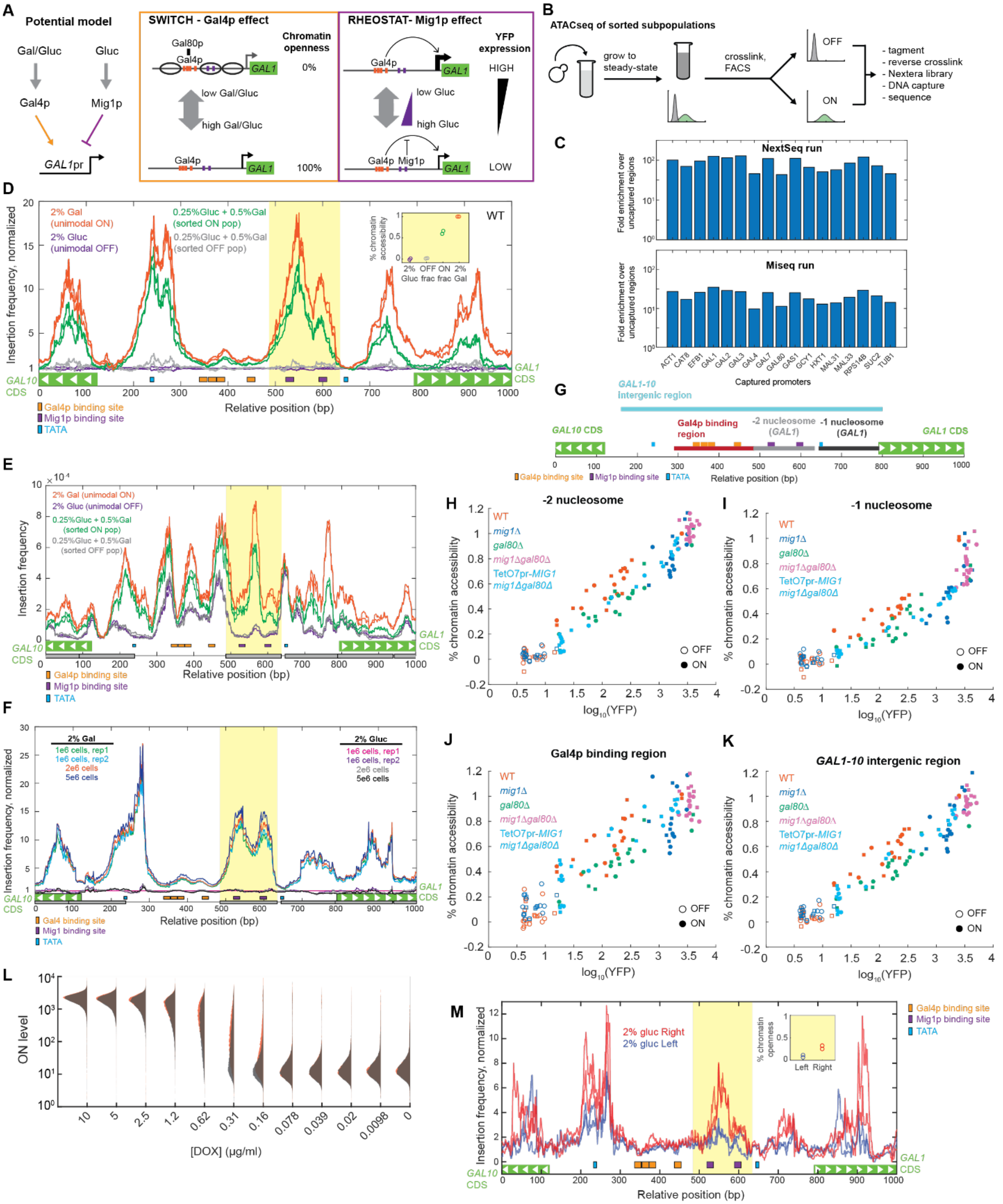

**Fig. S3.**

**Fig. S3. (Continued)**

Controls and additional supporting data for ATAC-seq experiments, related to Fig. 2. (A) Model for switch-and-rheostat decomposition at the *GAL1*pr based on the PHO system (Ref.7). Both Gal4p and Mig1p regulation integrate at the *GAL1* promoter (left): the switch from OFF to ON (box, left) is due to Gal4p-mediated remodeling of nucleosomes, while the rheostat (box, right) is through graded Mig1p binding to the open-chromatin conformation of the promoter. (B) Schematic of modified ATAC-seq protocol for assaying chromatin accessibility. Samples were crosslinked, sorted based on expression level, and then tagmented. To maximize the number of samples per sequencing run, a set of 17 promoters of interest were enriched by hybrid capture (Methods). (C) Fold enrichment in 17 promoters after hybrid capture for two different sequencing runs. (D) ATAC-seq insertion frequency across the *GAL1* promoter normalized to the average ATAC-seq signal in 2% glucose (purple, maximally repressed state - unimodal OFF) for wild-type cells grown in 2% galactose (orange, maximally induced state - unimodal ON) or a mixture of 0.25% glucose and 0.5% galactose (bimodal induction, cells were sorted into ON (green) and OFF (grey) subpopulations, as in (B)). The summed frequency over the genomic region occupied by the -2 nucleosome, highlighted in yellow, is used to calculate the percent chromatin openness metric (inset, Methods). (E) ATAC-seq insertion frequency of samples shown in Fig. S3D when not normalized to the 2% glucose condition. (F) Effect of number of cells input into the Nextera tagmentation (B) on ATAC-seq insertion frequency at the *GAL1* promoter (data from samples run on MiSeq). (G) Schematic illustrating four different windows in the *GAL1* promoter, where transposase insertions were summed to calculate percent chromatin openness summary metric. (H)-(K) Comparison of correlation between YFP level and chromatin openness when calculating openness metric by summing counts across the four different windows presented in (G); -2 nucleosome (H, Panel reproduced for comparison from Fig. 2A), -1 nucleosome (I), Gal4p binding region (J), and *GAL1-10* intergenic region (K). Circles represent replicate 1, squares represent replicate 2. (L) Histograms of *GAL1*pr-YFP expression level at different DOX concentrations (in a background of 2% glucose) for the *mig1Δgal80Δ* TetO7pr-*MIG1* reporter strain. (M) ATAC-seq insertion frequency at the *GAL1* promoter for the sorted left and right halves of the YFP expression distribution of *gal80Δ* cells grown in 2% glucose.

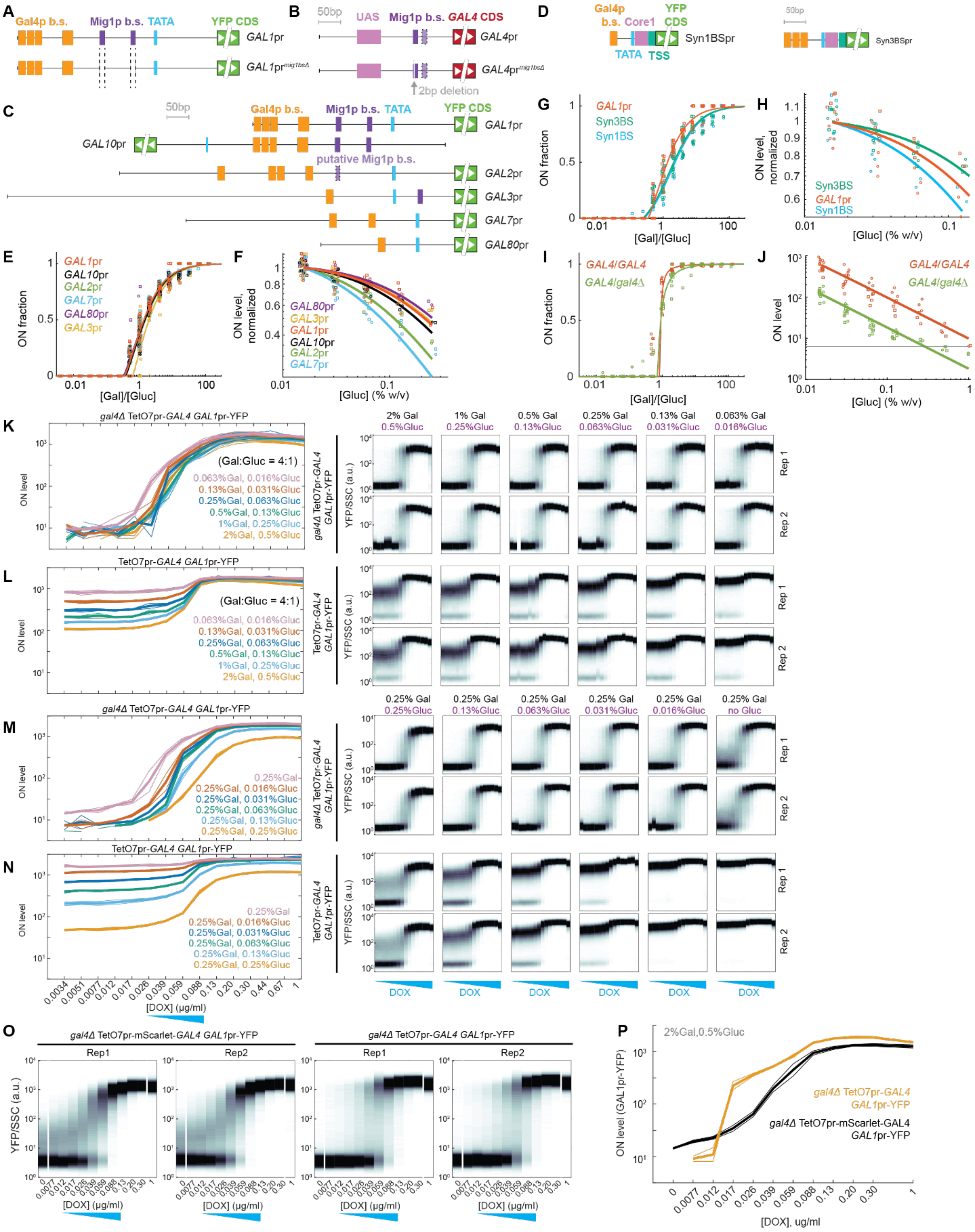

**Fig. S4.**

**Fig. S4. (Continued)**

Binding site distributions of reporter promoters and DOX titrations of Gal4p, related to Fig. 2. (A)-(D) Schematics of binding site distributions in the *GAL1*pr sequence before and after deletions to remove the Mig1p binding sites (A), *GAL4*pr sequence before and after the 2-bp deletion introduced to disrupt the functional Mig1p binding site and create the *GAL4*pr$^{mig1bs\Delta}$ sequence (B), promoters of the GAL gene reporter set (C), and synthetic promoter sequences (D). Putative Mig1p binding sites are by sequence prediction and have not yet been shown to be functional. (E)-(J) ON fraction vs. galactose:glucose ratio (E, G, and I) and ON level vs. glucose concentration (F, H, and J) for GAL gene promoters driving YFP (E-F), synthetic promoters with only Gal4p binding sites driving YFP (G-H), or *GAL1*pr-YFP for different mutants of the GAL pathway (I-J) at steady-state in a glucose-galactose double gradient (two replicates). X-values randomly jittered for visualization purposes. Grey line in (J) indicates the lower limit for being able to confidently distinguish an ON subpopulation from an OFF subpopulation. (K)-(N) DOX titration series of *gal4Δ* TetO7pr-*GAL4 GAL1*pr-YFP and TetO7pr-*GAL4 GAL1*pr-YFP strains along an axis of equal galactose:glucose ratio and varying absolute sugar concentrations (K and L respectively) or along a constant glucose axis with varying galactose (M and N respectively) (two replicates for each glucose-galactose condition) (raw DOX-titration density plots for each glucose-galactose condition on the right, and summarized ON level versus DOX concentration on the left). (O) DOX-titration density plots of YFP expression in *gal4Δ* TetO7pr-*GAL4 GAL1*pr-YFP and *gal4Δ* TetO7pr-mScarlet-*GAL4 GAL1*pr-YFP strains when grown in a 2% galactose + 0.5% glucose background (same experimental samples used for microscopy measurements shown in Fig. 2D). (P) *GAL1*pr-YFP expression level vs. DOX concentration of *gal4Δ* TetO7pr-*GAL4 GAL1*pr-YFP and *gal4Δ* TetO7pr-mScarlet-*GAL4 GAL1*pr-YFP strains at a fixed 2% galactose + 0.5% glucose concentration (two replicates).

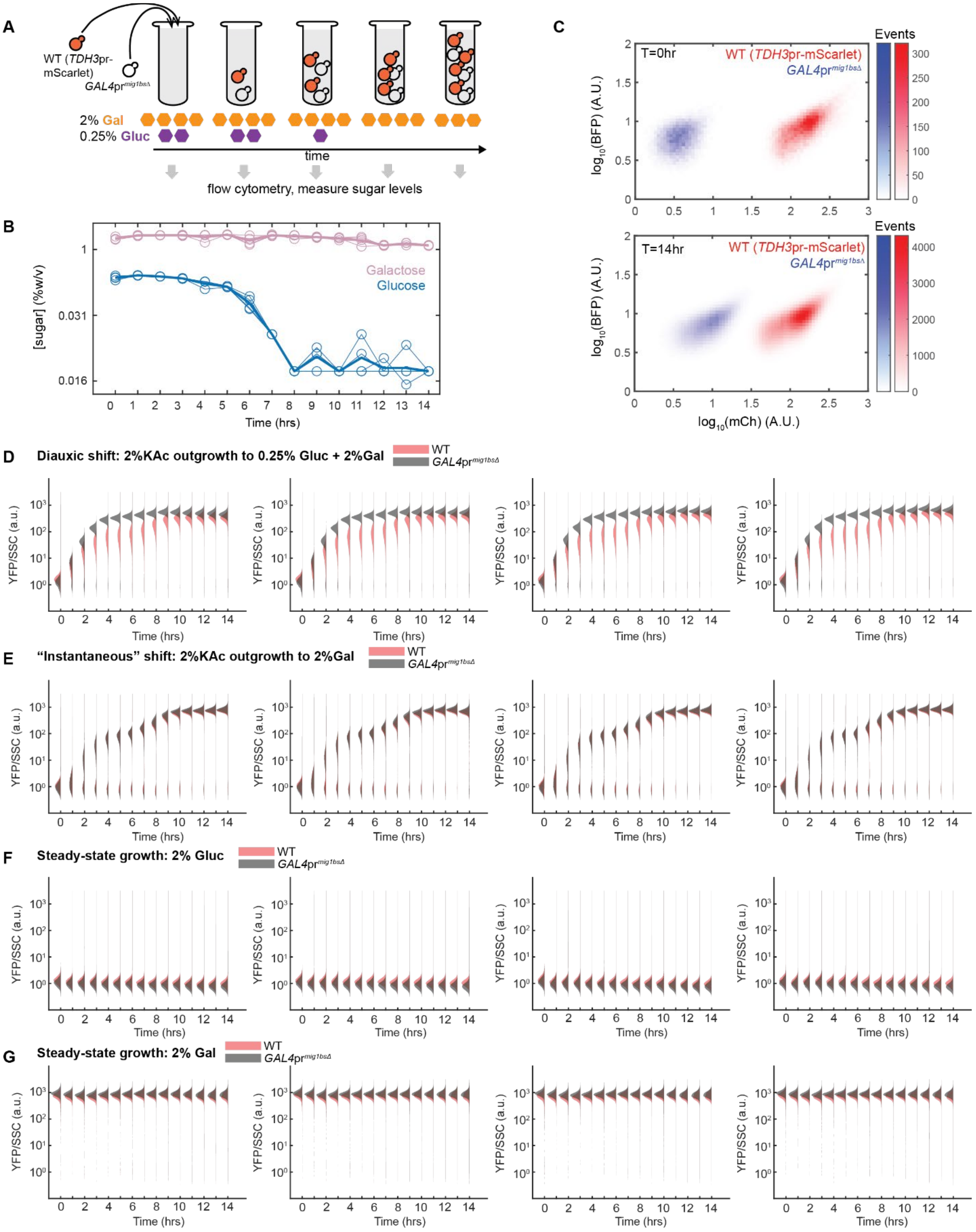
A
WT (TDH3pr-mScarlet)
GAL4pr mig1bsΔ
2% Gal
0.25% Gluc
time
flow cytometry, measure sugar levels
B
[sugar] (%w/v)
Galactose
Glucose
Time (hrs)
C
T=0hr
T=14hr
WT (TDH3pr-mScarlet)
GAL4pr mig1bsΔ
Events
log10(BFP) (A.U.)
log10(mCh) (A.U.)
D
Diauxic shift: 2%KAc outgrowth to 0.25% Gluc + 2%Gal
E
"Instantaneous" shift: 2%KAc outgrowth to 2%Gal
F
Steady-state growth: 2% Gluc
G
Steady-state growth: 2% Gal
WT
GAL4pr mig1bsΔ
YFP/SSC (a.u.)
Time (hrs)

**Fig. S5.**

**Fig. S5. (Continued)**

Co-culture strain segmentation, sugar depletion, and *GAL1*pr-YFP induction time-courses for fitness competitions between wildtype and *GAL4*pr$^{mig1bs\Delta}$ strains, related to Fig. 4. (A) Overview of the experimental procedure for time series measurements of the competitive fitness of a wild-type strain and a *GAL4*pr$^{mig1bs\Delta}$ strain during a diauxic shift, in parallel with measurements of their *GAL1*pr-YFP expression and sugar concentrations. For segmentation of the strains in co-culture, the wildtype strain constitutively expressed mScarlet. (B) Sugar depletion measurements, starting from a mixture of 2% galactose and 0.25% glucose, during the diauxic shift competition shown in Fig.4A (thick lines represent averages over four replicates in thin lines), as measured using an enzymatic assay and absorbance at 340nm (A340) (Methods). (C) Segmentation of wild type (constitutively expressing mScarlet) and *GAL4*pr$^{mig1bs\Delta}$ based on red fluorescence at both the beginning (t=0) and end (t=14hr) of the diauxic shift time-course. (D)-(G) Histograms of *GAL1*pr-YFP expression over a 14-hr period for the diauxic shift (D), “instantaneous” shift (E), steady-state glucose growth (F), and steady-state galactose growth (G) competition experiments shown in Fig. 4A.

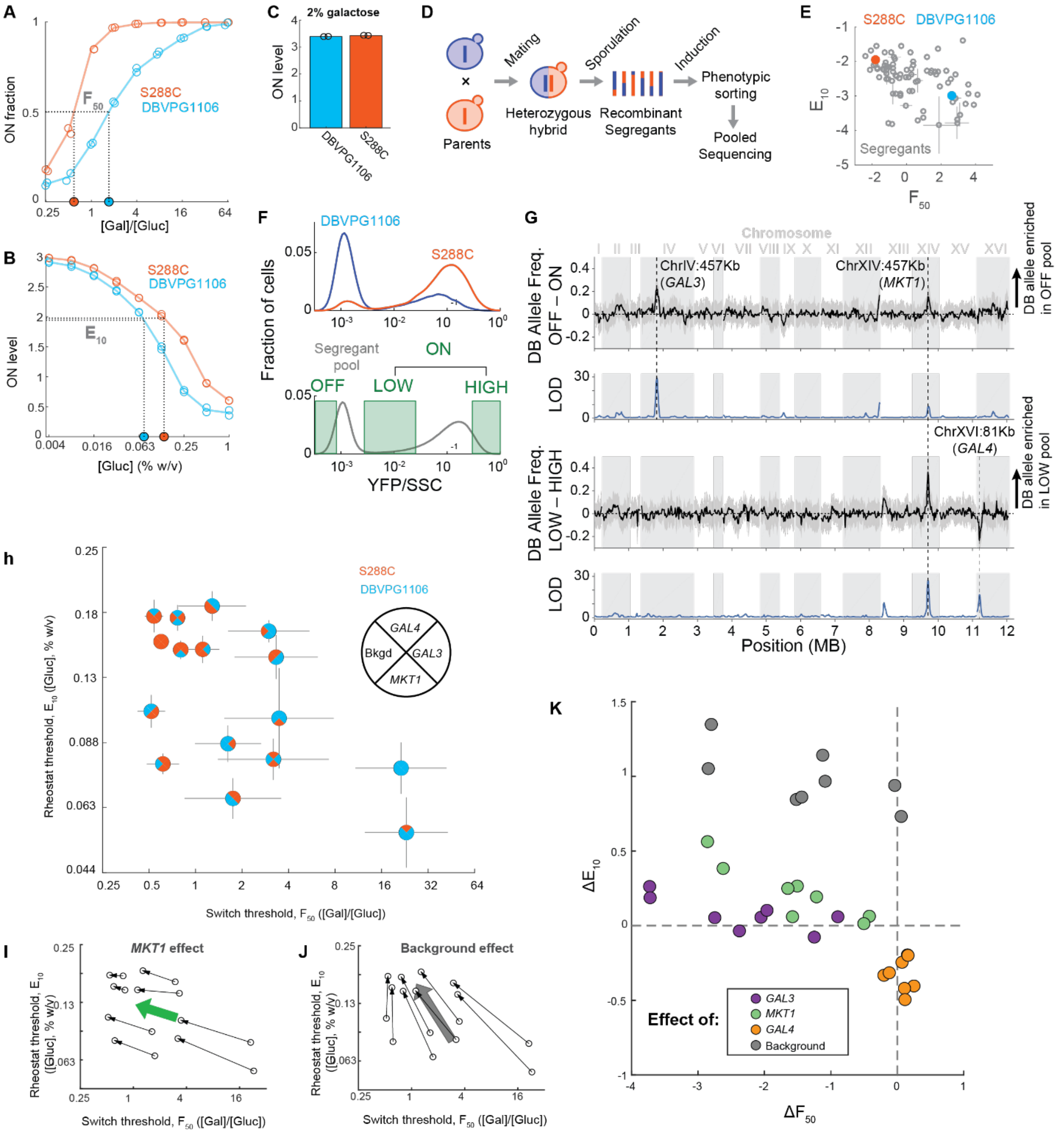


### Fig. S6.

Bulk segregant analysis and combinatorial effects of strain background and GAL3, MKT1, and GAL4 alleles, related to Fig. 4. (A)-(B) Metrics for decision (A) and level (B) “setpoints” in S288C (blue) and DBVPG1106 (red) strain backgrounds grown in a glucose gradient in a background of a 0.25% galactose concentration. The lines connect the averages over six replicates at each glucose concentration. (A) The switch threshold ($F_{50}$) corresponds to the galactose:glucose ratio at which 50% of cells are induced (ON fraction is 50%). (B) The rheostat threshold ($E_{10}$) corresponds to the glucose concentration at which expression level is 10% of maximal expression in 2% galactose. (C) ON level of DBVPG1106 and S288C parent strains when grown in 2% galactose medium. (D) Schematic of bulk segregant analysis strategy. (E) $E_{10}$ versus $F_{50}$ across 90 haploid segregants of the DBVPG1106 x S288C cross. Parent phenotypes are shown as filled circles: DBVPG1106 (blue), S288C (red). (F) *GAL1*pr-YFP histograms of parent strains DBVPG1106 (blue) and S288C (red) (top) and a pool of haploid segregants (gray, bottom) in the sorting conditions, 0.25% glucose + 0.25% galactose. Green boxes are a schematic of the gates used to sort segregant cells into 3 phenotype pools (OFF, LOW, and HIGH) for sequencing (gates used in actual sorting experiment are shown in Fig. S7C). ON pool allele counts are a computational sum of the LOW and HIGH pool allele counts (Methods). (G) Genome-wide plots of differential allele frequency and log-odds-ratio (LOD) as computed by the MULTIPOOL algorithm (Methods). Top two plots show the OFF/ON comparison; bottom two plots show the LOW/HIGH comparison. Positive values represent enrichment in DBVPG1106 (DB) strain background. (H) Scatterplot of $E_{10}$ versus $F_{50}$ from combinatorial allele swaps of *GAL3*, *GAL4*, and *MKT1* between the S288C (red quadrant) or DBVPG1106 (blue quadrant) strain backgrounds. (I)-(J) $E_{10}$ versus $F_{50}$ for all 16 combinations of S288C ("S") or DBVPG1106 ("D") strain background, *GAL3* allele, *MKT1* allele, and *GAL4* allele (similar to Fig. 4B-C). Effects of switching *MKT1* allele (I) or strain background (J) from DBVPG1106 to S288C while holding other genetic variables constant are shown as black arrows, with thick arrows representing the average effect. (K) The effects shown in (I)-(J) and Fig. 4B-C are plotted as differences in $E_{10}$ versus differences in $F_{50}$.

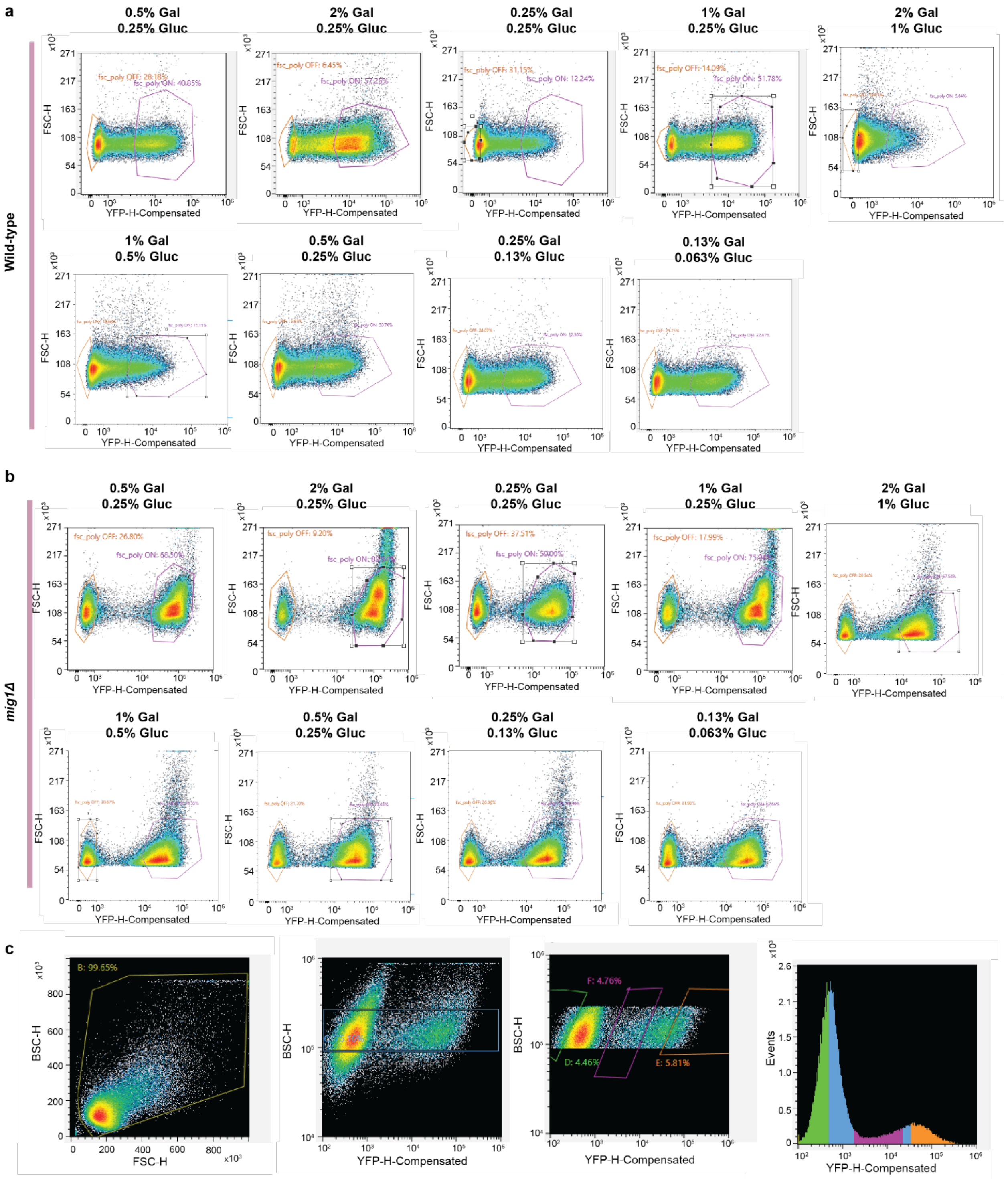


**Fig. S7.**

FACS sorting gates used for ATAC-seq sample preparation, and bulk segregant analysis. (A)-(B) Polygonal gates, in forward scatter (FSC) versus FITC space, used for sorting bimodal ATAC-seq samples of wildtype (A) and *mig1Δ* (B) strain backgrounds. (C) Polygonal gates used for sorting intercrossed segregants as illustrated in Fig. S6f.

**Table S1-S3:** see “Table_S1.xlsx”, “Table_S2.xlsx”, “Table_S3.xlsx”.

**Table S4.**

| **Strain** | **Fit** | **Equation** | **Parameters** | **Appears in fig.** |
|---|---|---|---|---|
| Wildtype | ON fraction | $f_{on} = \begin{cases} 0 & , r < A \\ \frac{r-A}{r-A+K_f}, & r \geq A \end{cases}$ | $A = 0.52 \pm 0.08$<br>$K_f = 0.77 \pm 0.08$<br>$fit\ s.e. = 0.04$ | Fig. 1J |
| *mig1Δ* | ON fraction | $f_{on} = \begin{cases} 0 & , r < A \\ \frac{r-A}{r-A+K_f}, & r \geq A \end{cases}$ | $A = 0.17 \pm 0.05$<br>$K_f = 0.6 \pm 0.1$<br>$fit\ s.e. = 0.1$ | Fig. 1J |
| *gal80Δ* | ON fraction | $f_{on} = A_f \log(r) + B_f$ | $A_f = 0.004 \pm 0.001$<br>$B_f = 0.969 \pm 0.003$<br>$fit\ s.e. = 0.02$ | g. 1J |
| *mig1Δ gal80Δ* | ON fraction | $f_{on} = A_f \log(r) + B_f$ | $A_f = 0$<br>$B_f = 1$<br>$fit\ s.e. = 7 \cdot 10^{-10}$ | Fig. 1J |
| Wildtype | ON level | $\mu_{on} = \mu_{max} \frac{K_\mu}{[Gluc]+K_\mu}$ | $\mu_{max} = 4 \cdot 10^3 \pm 1 \cdot 10^3$<br>$K_\mu = 0.019 \pm 0.008$<br>$fit\ s.e. = 2 \cdot 10^2$ | Fig. 1K |
| *mig1Δ* | ON level | $\mu_{on} = \mu_{max} \frac{K_\mu}{[Gluc]+K_\mu}$ | $\mu_{max} = 2.7 \cdot 10^3 \pm 0.1 \cdot 10^3$<br>$K_\mu = 4 \pm 6$<br>$fit\ s.e. = 3 \cdot 10^2$ | Fig. 1K |
| *gal80Δ* | ON level | $\mu_{on} = \mu_{max} \frac{K_\mu}{[Gluc]+K_\mu}$ | $\mu_{max} = 9 \cdot 10^3 \pm 2 \cdot 10^3$<br>$K_\mu = 0.008 \pm 0.003$<br>$fit\ s.e. = 3 \cdot 10^2$ | Fig. 1K |
| *mig1Δ gal80Δ* | ON level | $\mu_{on} = \mu_{max} \frac{K_\mu}{[Gluc]+K_\mu}$ | $\mu_{max} = 4.2 \cdot 10^3 \pm 0.2 \cdot 10^3$<br>$K_\mu = 1.6 \pm 0.6$<br>$fit\ s.e. = 6 \cdot 10^2$ | Fig. 1K |
| Wildtype | ON fraction | $f_{on} = \frac{r^n}{r^n + K_f^n}$ | $K_f = 1.45 \pm 0.07$<br>$n = 1.9 \pm 0.1$<br>$fit\ s.e. = 0.06$ | Fig. S2B |
| Wildtype | ON fraction | $f_{on} = \frac{r}{r + K_f}$ | $K_f = 1.6 \pm 0.2$<br>$fit\ s.e. = 0.10$ | Fig. S2B |

Fitting Values.
